# FANCD2 and RAD51 recombinase directly inhibit DNA2 nuclease at stalled replication forks and FANCD2 acts as a novel RAD51 mediator in strand exchange to promote genome stability

**DOI:** 10.1101/2021.07.08.450798

**Authors:** Wenpeng Liu, Piotr Polaczek, Ivan Roubal, Yuan Meng, Won-chae Choe, Marie-Christine Caron, Carl A. Sedgeman, Yu Xi, Changwei Liu, Qiong Wu, Li Zheng, Jean-Yves Masson, Binghui Shen, Judith L. Campbell

## Abstract

FANCD2 protein, a key coordinator and effector of the interstrand crosslink repair pathway, is also required to prevent excessive nascent strand degradation at hydroxyurea-induced stalled forks. The RAD51 recombinase has also been implicated in regulation of resection at stalled replication forks. The mechanistic contributions of these proteins to fork protection are not well understood. Here, we used purified FANCD2 and RAD51 to study how each protein regulates DNA resection at stalled forks. We showed that FANCD2 inhibits fork degradation in two ways: 1) the N-terminal domain of FANCD2 inhibits DNA2 nuclease activity by directly binding to DNA2. 2) independent of dimerization with FANCI, FANCD2 itself stabilizes RAD51 filaments to inhibit multiple nucleases, including DNA2, MRE11, and EXO1. Unexpectedly, we uncovered a new FANCD2 function: by stabilizing RAD51 filaments, FANCD2 acts as a “RAD51 modulator” to stimulate the strand exchange activity of RAD51. Our work biochemically explains non-canonical mechanisms by which FANCD2 and RAD51 protect stalled forks. We propose a model in which the strand exchange activity of FANCD2 provides a simple molecular explanation for genetic interactions between FANCD2 and the BRCA2 mediator in the FA/BRCA pathway of fork protection

## Introduction

Successful completion of DNA replication requires the integration of many proteins and pathways that protect, repair and/or restart replication forks. The principles underlying how these pathways interact and are regulated to maximize genome stability have yet to be determined. Fanconi anemia is a rare disease of bone marrow failure, developmental abnormalities, and cancer predisposition. At the cellular level it is diagnosed by sensitivity to DNA ICLs (interstrand crosslinking agents) and genome instability. Fanconi anemia is a multigenic disease defined by at least 22 complementation groups, including many regulatory components, nucleolytic activities, and HDR (homology directed repair) genes. The component genes suggest a coherent pathway for maintaining genome stability during DNA replication that goes beyond ICL repair and includes response to many additional types of replication stress (Boisvert and Howlett, 2014; Federico et al., 2018). The multigenic character of the FA pathway lends itself to comprehensive genetic and biochemical dissection (Carr and Lambert, 2013; Chaudhury et al., 2013; Chaudhury et al., 2014; Hashimoto et al., 2010; Kottemann and Smogorzewska, 2013; Lossaint et al., 2013; Petermann et al., 2010; Raghunandan et al., 2015; Schlacher et al., 2011; Schlacher et al., 2012; Sobeck et al., 2006; Yeo et al., 2014).

FANCD2 is a key regulator of the FA pathway and the focus of our current studies (Kottemann and Smogorzewska, 2013; Timmers et al., 2001). During canonical replication-coupled repair of ICLs, after a replication fork encounters an ICL, FANCD2 and a related protein FANCI, are phosphorylated by activated ATR kinase. A FANCD2/FANCI heterodimer is also formed, and FANCD2 in this heterodimer, but not free FANCD2, is mono-ubiquitylated by the FA core complex, containing nine FA proteins, including the FANCL E3 ligase complex and several associated proteins. FANCD2-ubi is involved in both activation of repair events and also is directly required in the later enzymatic repair steps at strand breaks (Knipscheer et al., 2009; Kottemann and Smogorzewska, 2013; Long et al., 2014; Long et al., 2011; Long and Walter, 2012; Raschle et al., 2008). The role of ubiquitin is to enforce stable binding of FANCD2/FANCI to DNA, specifically by clamping FANCD2-ubi/FANCI heterodimers onto DNA for DNA repair (Alcon et al., 2020; Rennie et al., 2020; Tan et al., 2020; Wang et al., 2020).

FANCD2 also appears to have both ubiquitin-independent and FANCI-independent functions. In addition to its role in ICL repair, FANCD2 is required in the BRCA2-regulated, RAD51-mediated replication fork protection pathway, irrespective of the source of DNA damage (Chaudhury et al., 2013; Raghunandan et al., 2015; Schlacher et al., 2011; Schlacher et al., 2012). While several studies of fork protection implied that non-ubiquitylatable FANCD2 (FANCD2-K561R) could not restore fork protection to patient-derived FANCD2-defective cells (Kais et al., 2016; Schlacher et al., 2012), other results support that FANCD2 is likely to have constitutive functions, at least for low levels of replication stress, such as endogenous stress (Boisvert and Howlett, 2014; Federico et al., 2018). With respect to ubiquitylation, study of FANCD2 knockout and knock-in cell lines showed that cells expressing only non-ubiquitylatable FANCD2-K561R had much less severe phenotypes than cells with a FANCD2 knockout (Tian et al., 2017). Complementary studies showed that mutants defective in the trans-acting FA core complex components responsible for ubiquitylation of FANCD2 are less sensitive to replication fork stalling agents than FANCD2 knockdowns or knockouts (Thompson et al., 2017). Importantly, one of us reported that FANCD2 can protect forks by different mechanisms than FANCA/C/G, members of the core complex (Liu et al., 2020). FANCD2 has been shown to interact with RAD51, a key player/regulator in fork protection, and to do so in a ubiquitylation-independent, but HU-stimulated manner (Chen et al., 2017). FANCD2 also has FANCI independent functions (Chaudhury et al., 2013; Dubois et al., 2019; Thompson et al., 2017). FANCD2 deficient cells are HU and aphidicolin (a DNA polymerase inhibitor) sensitive, while FANCI cells are not (Thompson et al., 2017). These results stimulated our interest in studies of ubiquitin- and FANCI-independent roles of FANCD2.

Fork protection implies protection from nucleases. Several nucleases have been implicated in nascent DNA degradation at stalled forks, in both fork-protection proficient and fork-protection deficient cells (Schlacher et al., 2011; Schlacher et al., 2012; Thangavel et al., 2015). Prominent among these is the DNA2 helicase/nuclease, which seems to play a role in several different fork protection subpathways (Liu et al., 2020; Rickman et al., 2020). DNA2 is essential for replication in normal yeast cells (Zheng et al., 2020). Multi-tasking DNA2 removes long 5’ ssDNA flaps in the presence of Pif1 helicase during non-canonical Okazaki fragment processing (Budd et al., 2006; Diffley, 2020; Zheng et al., 2020); performs long-range resection of DSBs to provide 3’ ends for BRCA2-mediated, RAD51 filament formation and strand invasion during HDR; and prevents deleterious fork reversal in yeast (Hu et al., 2012). At stalled forks, human DNA2 is required for repair of stalled replication forks, where it mediates limited resection, putatively on a reversed fork, to promote fork restart, in replication fork protection proficient cells (Thangavel et al., 2015). We discovered that DNA2 deficient cells are sensitive to inter- or intra-strand crosslinks induced by cisplatin or formaldehyde. Paradoxically, depletion of DNA2 in cells deficient in FANCD2 rescued ICL sensitivity in FANCD2 mutants, suggesting that DNA2 became toxic in the absence of fork protection (Karanja et al., 2014; Karanja et al., 2012). We and others provided direct evidence that in cells defective in fork protection, DNA2-mediated over-resection occurs at replication forks stalled by exogenous DNA damaging agents or replication inhibition and that FANCD2 is likely to regulate this resection (Higgs et al., 2015; Higgs et al., 2018; Jiang et al., 2015; Liu et al., 2020; Liu et al., 2016; Rickman et al., 2020; Thangavel et al., 2015; Tian et al., 2017; Wang et al., 2015). The question remains as to how FANCD2 regulates DNA2, however.

RAD51 recombinase can also be inhibitory to over-resection.by DNA2 inhibition in vivo (Thangavel et al., 2015). A dominant negative RAD51 mutant leads to excessive degradation of nascent DNA in a RAD51T131P/RAD51 heterozygote (Wang et al., 2015). FANCD2 and RAD51 are epistatically linked in fork protection (Schlacher et al., 2012). In vivo, however, it is not known if RAD51 and FANCD2 act independently or together.

Recently, several studies have suggested that FANCD2 and BRCA2, the RAD51 mediator, perform parallel or compensatory functions in fork protection and fork recovery after collapse (Kais et al., 2016; Lachaud and Rouse, 2016; Michl et al., 2016). Since BRCA2 is thought to stabilize RAD51 filaments, we hypothesized that FANCD2 may provide a backup source of this BRCA2 function in response to replication stress. This mechanism is supported by the fact FANCD2 alone and FANCD2/FANCI heterodimers interact physically with RAD51 (Berti et al., 2013; Chen et al., 2017; Sato et al., 2016; Thangavel et al., 2015; Thangavel et al., 2010). FANCD2/FANCI complexes have been shown to increase RAD51 levels on DNA, but the specific contribution of FANCD2 itself and the relationship of this observation to suppression of BRCA2^−/−^ defects has not been established.

In this work we delved deeper into the mechanisms of FANCD2 and RAD51 fork protection by comparing the effect of FANCD2 deficiency on replication forks stalled due to replication stress induced by different damaging agents. Our in vivo results confirm that FANCD2 is required to protect stalled replication forks from DNA2-dependent over-resection after acute stress. We identified at least two potential mechanisms by which FANCD2 protects nascent DNA from nucleolytic resection in vitro: 1) FANCD2 inhibits DNA2 nuclease activity and binds directly to the DNA2 nuclease domain; and 2) FANCD2 stabilizes RAD51 ssDNA filaments which prevent nucleolytic digestion by multiple nucleases. Surprisingly, we also find that FANCD2, like BRCA2, acts as a mediator in RAD51-dependent strand exchange. FANCD2 promotes RAD51-mediated strand exchange activity by stabilizing RAD51 on ssDNA, but not, like BRCA2 by inhibiting the RAD51 ATPase or RAD51 dsDNA binding activities, and not requiring FANCD2 DNA binding motifs. The ability to stimulate strand exchange provides a novel mechanistic explanation for the well-documented dependency of BRCA2^−/−^ tumors on FANCD2, the suppression of BRCA1/2^−/−^ phenotypes by elevated levels of FANCD2 (Kais et al., 2016; Lachaud and Rouse, 2016; Michl et al., 2016), and adds a new dimension to how FANCD2 deficiency leads to inhibition of fork protection and to genome instability (Schlacher et al., 2012).

## Results

### FANCD2 is Required for Replication Fork Protection after Acute Replication Stress

Multiple pathways are involved in stalled replication fork repair, and forks undergo progressive changes in architecture during chronic stalling (Helleday et al., 2008; Lemacon et al., 2017; Petermann et al., 2010; Quinet et al., 2017). Previous studies showed that FANCD2 can protect nascent DNA from degradation upon the replication stress. However, the major question is how FANCD2 protects the stalled fork and what structure it is being acted upon (Petermann et al., 2010; Piberger et al., 2019). To investigate this question, we first verified over-resection in FANCD2 deficient cells with RPA2 phosphorylation as a surrogate for measuring ssDNA arising during resection in the presence of HU. Cells were treated with HU for 0-8 h and nuclear extracts prepared. At all-time points, we observe drastically increased RPA-p levels (resection) in nuclear extracts of HU-treated PD20 FANCD2^−/−^ deficient cells compared to PD20:FANCD2 (FANCD2-complemented) cells, where there was very little resection (Figure 1A). This is consistent with other published results showing that FANCD2 protects stalled forks from nascent DNA degradation.

**Figure 1.**
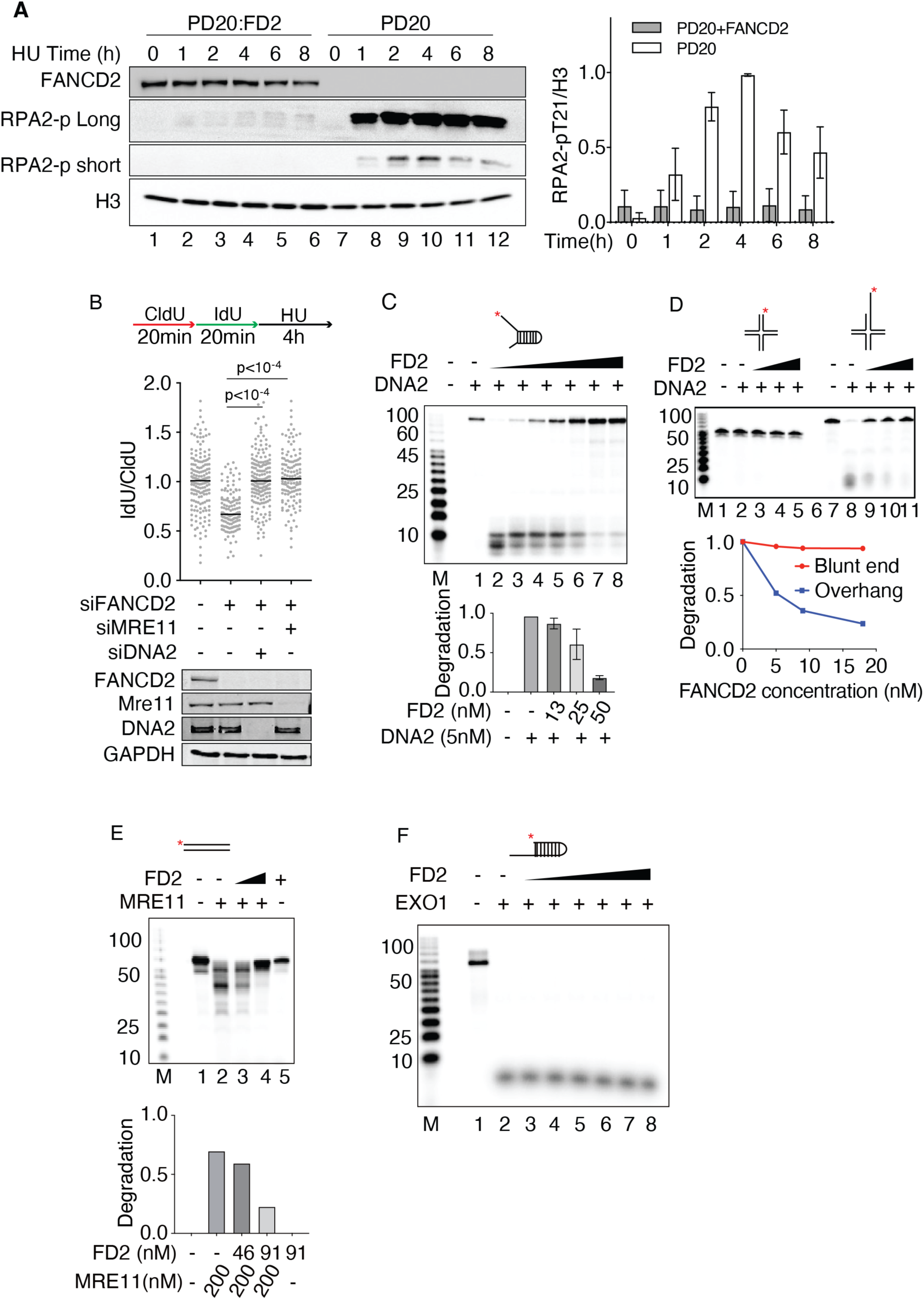
FANCD2 prevents DNA2 and MRE11 mediated nascent DNA degradation. (A) FANCD2 prevents resection in response to HU. PD20 cells and PD20:FANCD2 complemented cells treated with 3 mM hydroxyurea for 0-8 hours. HU was added and samples were taken at 0, 2, 4, 6, and 8 h. Nuclear extract was prepared, as illustrated by the H3 loading control, and analyzed for resection by western blot. “RPA2-p long” refers to long exposure time monitoring RPA2 T21 phosphorylation; “RPA2-short” is a short exposure time monitoring RPA2 T21 phosphorylation. The RPA2-p and histone H3 blot intensity were measured with ImageJ. Then RPA2-p value was normalized to histone H3. The graph value is the average of two experiments; error bar is standard deviation. (B) Over-resection of nascent DNA in HU-treated FANCD2 deficient cells is reduced by either DNA2 or MRE11 depletion. U2OS cells were co-transfected with 12 nM siRNA for each indicated gene. 72 hours post-transfection, as indicated on the top of the panel, cells were pulsed by CldU and IdU, followed by 4 mM HU for 4h. The cells were harvested and analyzed by fiber spreading assay. The IdU and CldU tract lengths were measured, and the ratio was graphed (≥150 fibers were analyzed). One-way ANOVA test was performed, n=2. Western blots show the quantification of knockdown. (C) DNA2 nuclease activity is inhibited by FANCD2-His on a single-stranded fork. Increasing amounts of FANCD2-His were preincubated in DNA2 nuclease reaction (8 μl) mix for 30 min at 4°C. DNA substrate (87 FORK, 15 nM, 1 μL) was added, and the reaction was incubated for 30 min at 37°C. All reactions contained equal amounts of FANCD2 diluent. Lane 1, DNA alone; lane 2, DNA2 alone; lanes 3-8, DNA2 (1 nM) plus 13, 25, and 50 nM FANCD2 in duplicate, respectively. (D) Inhibition of DNA2 by FANCD2 on reversed forks. Reactions were as in panel C. Lane 1 and 7, DNA alone; lane 2 and 8, DNA2 alone; lanes 3-5 and lanes 9-11, DNA2 (1 nM) plus 5, 9, and 18 nM FANCD2, respectively. Verification of the reversed fork structures is presented in Figure S1F. In lieu of repeating assays with one concentration of FANCD2 on the additional substrates, significance was indicated by showing concentration dependence, which is more informative. (E) FANCD2-His inhibits MRE11 on a dsDNA substrate. FANCD2-His was preincubated in MRE11 nuclease reaction mix on ice for 30 min. Then substrate added to activate the reaction in 37 °C for 30mins. dsDNA substrate is shown in the top of the panel. Quantification shows the degradation levels. (F) FANCD2 does not inhibit EXO1 on an overhang substrate. FANCD2 was added as indicated, FANCD2-His was preincubated in EXO1 nuclease reaction mix on ice for 30 min. Then substrate added to activate the reaction in 37 °C for 30mins.

We next asked if knockdown of DNA2 can rescue the over-resection observed after 4 h of HU treatment, in keeping with the fact that the cisplatin and formaldehyde sensitivity of the FANCD2^−/−^ PD20 patient cell line can be suppressed by DNA2 knockdown (Karanja et al., 2014). Using single molecule tracking of nascent DNA before and after brief (4h) HU treatments, as reported previously, we see over-resection in FANCD2 depleted cells (Figure 1B), supporting use of RPA-p as the readout for resection shown in Figure 1A. We then confirm that degradation of nascent DNA in FANCD2-deficient cells upon HU treatment can be rescued by knockdown of DNA2 and also by knockdown of MRE11 nuclease (Figure 1B). This is consistent with the previous studies. We also verified that over-resection did not require fork reversal by SMARCAL1 or ZRANB3 nor cleavage by MUS81/SLX4 (not shown), as one of us had previously demonstrated (Liu et al., 2020). We conclude that MRE11 and DNA2 may function as alternative nucleases or function sequentially in stalled fork processing, as they do at DSBs (Symington and Gautier, 2011).

### FANCD2 Inhibits DNA2 Nuclease Activity in vitro Providing a Mechanism for FANCD2’s in vivo Role in Fork Protection

Since over-resection in PD20 cells treated with HU is massively increased over the FANCD2-complemented cells (Figure 1A), and since FANCD2 and DNA2 have been shown to interact in vivo in a DNA-independent fashion, suggesting a protein/protein interaction (Karanja et al., 2012), we tested whether FANCD2 directly regulates degradation by DNA2 nuclease. FANCD2-His was purified from SF9 insect cells (Figure S1A) (Roques et al., 2009) and was shown to bind dsDNA (Figure S1B) and to be free of nuclease activities under the conditions used here (Figure S1C). When FANCD2 was added to a DNA2 nuclease reaction containing a forked substrate, significant inhibition of DNA2 nuclease was observed, even in the presence of high levels (5 nM) of DNA2 (Figure 1C). Inhibition is likely due to FANCD2 protein and not reaction conditions since all reactions contained the same amount of FANCD2 diluent. In these experiments, the substrate partially mimics a stalled replication fork with single-stranded DNA arms at the dsDNA junction. DNA2 processes substrates with several different configurations, such as unligated 5’ flaps on Okazaki fragments or on base excision repair intermediates, single-stranded DNA, or 5’ overhangs on regressed replication forks during replication fork stress/stalling or DSB resection during homologous recombination. As shown in Figures 1D, S1D and S1E, FANCD2 inhibits DNA2 nuclease on each of these structures. (Sequences of all oligonucleotide substrates used are provided in Table S2.) The substrates mimicking reversed fork DNAs with or without a 5’ overhang (Figure 1D) were generated and validated as described in Figure S1F.

FANCD2 forms stable complexes with FANCI in vivo and in vitro, although only 20% of the FANCD2 in the cell co-IPs with FANCI (Alcon et al., 2020; Tan et al., 2020). We next tested if FANCI also inhibits DNA2 nuclease activity. We showed that there was no inhibition of DNA2 nuclease by FANCI alone (Figure S1G), and that addition FANCI together with FANCD2 did not further inhibit the DNA2 nuclease activity (Figure S1H).

Since FANCD2 also prevents nascent strand degradation by MRE11, we next addressed if FANCD2 inhibits MRE11 (Figure 1E). We found that MRE11 activity on duplex DNA was inhibited by FANCD2, also consistent with a protein/protein interaction (Roques et al., 2009). We also tested the effect of FANCD2 on EXO1 (Figure 1F). EXO1 was not inhibited by FANCD2, so there is specificity to the inhibition of MRE11 and DNA2 by FANCD2.

### How Does FANCD2 Inhibit DNA2 Nuclease?

Since FANCD2 binds to DNA (Niraj et al., 2017) as well as binding to DNA2, we next asked whether inhibition was mediated by a FANCD2 protein/DNA interaction and/or DNA2/FANCD2 protein/protein interaction. To investigate whether DNA binding by FANCD2 was involved in the inhibition of DNA2, we used a FANCD2-F1+F3Mut, which is defective, though not completely blocked, in DNA binding (Figure S2A and S2B) (Niraj et al., 2017). Like WT FANCD2, FANCD2-F1+F3Mut showed no nuclease activity itself (Figure 2A, controls, left) and strongly inhibited DNA2 nuclease both on the fork structure (Figure 2A right) and on the reversed fork structure (Figure 2B), although it bound to ssDNA 10-fold less efficiently than WT FANCD2 (Figure S2B), consistent with the previous characterization of the FANCD2-F1+F3Mut protein (Niraj et al., 2017). This suggests that inhibition does not occur by simply blocking the DNA substrate or competing with DNA2 for the substrate and suggests that direct protein/protein interaction may account for DNA2 inhibition.

**Figure 2.**
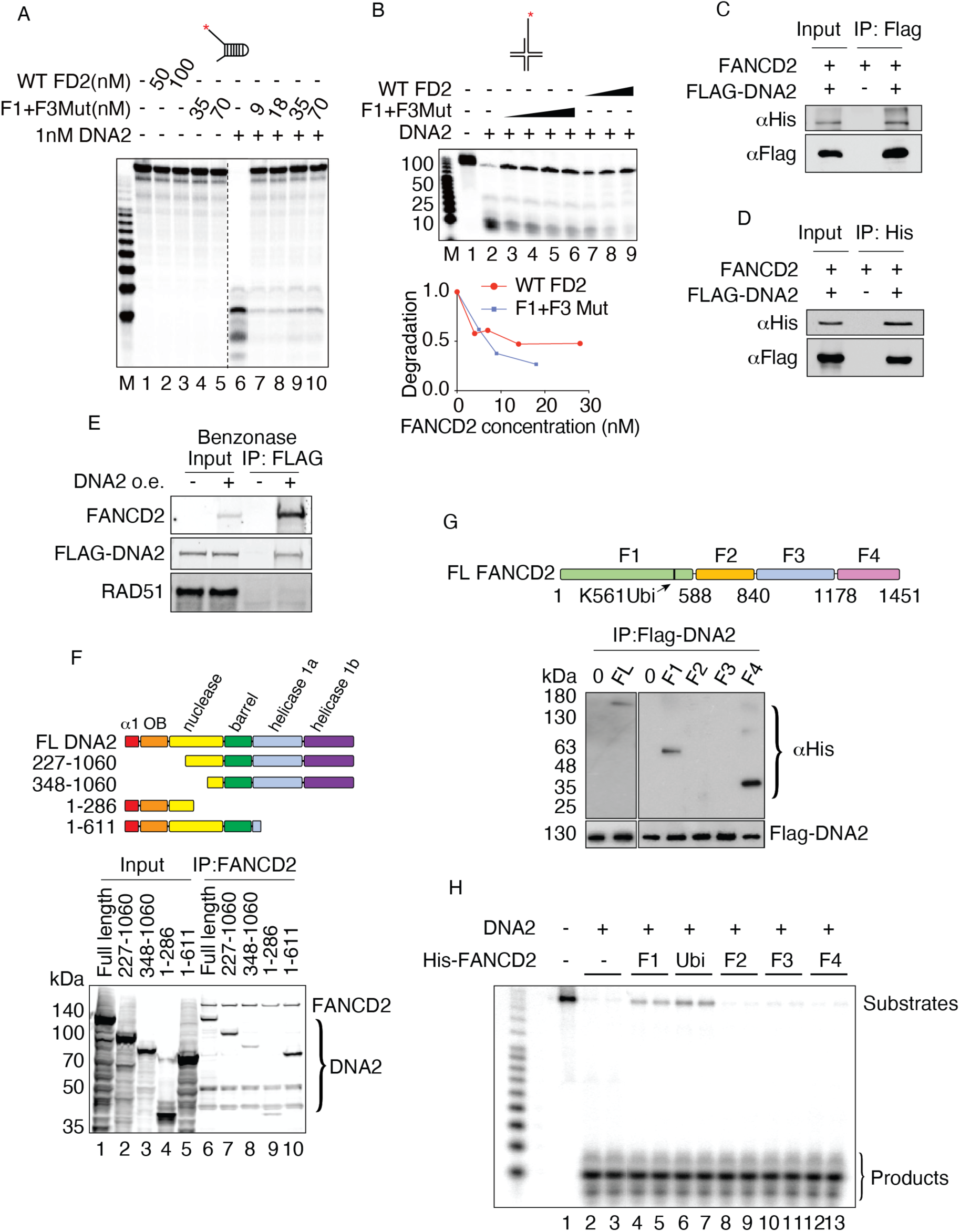
FANCD2/DNA interaction and FANCD2/DNA2 contribute to DNA2 nuclease inhibition. (A) Lack of Inhibition of hDNA2 by the hFANCD2-F1+F3Mut DNA binding mutant protein – 87 FORK substrate. Conditions were as in Figure 1 except that mutant FANCD2 was used. Lanes 1-5, FANCD2 wild type (WT) and F1+F3Mut alone, lane 6, DNA2 (1 nM) alone; lanes 7-10, DNA2 (1 nM) plus 9, 18 35, and 70 nM FANCD2-F1+F3Mut protein, respectively. (B) Lack of inhibition of hDNA2 by the hFANCD2-F1+F3Mut DNA binding mutant protein – reversed fork substrate. Quantification at the bottom shows the degradation levels. (C) FANCD2 and DNA2 interact in vitro. FLAG-DNA2 was over expressed in 293T cells, and purified by binding to M2 FLAG beads. The beads were washed with lysis buffer and then incubated with purified FANCD2 protein (1 μg/ml) at 4°C for 1 h. Beads were washed with PBS 5 times and then in 2XSDS loading buffer followed by western blotting. Empty M2 beads incubated with FANCD2 served as negative control. (D) FLAG-DNA2 is immuno-precipitated by His-tagged FANCD2 in vitro. 1 μg/ml DNA2 protein was incubated with FANCD2-His bound to Ni-NTA beads at 4°C for 1 hour. Beads were washed 5 times with PBS, and then boiled in 2XSDS loading buffer for western blotting. Empty Ni-NTA beads were incubated with FLAG-DNA2 as negative control. (E) DNA2 and FANCD2 interact in the absence of DNA. FLAG-DNA2 was over expressed in wild type U2OS cells, and nuclear extract was prepared; Benzonase was added to remove DNA. FLAG-DNA2 was pulled down with M2-beads. The beads were washed with nuclear preparation buffer and eluted with FLAG peptide. The elution was prepared for immunoblot. See Method Details. (F) FANCD2 interacts with the N terminal domain of DNA2. (Top) Map of DNA2 domains and truncations of DNA2. (Bottom) Coimmunoprecipitation of the full length (FL) DNA2 and truncated DNA2 proteins using FANCD2 antibody. Cell lysates with overexpressed FLAG-DNA2 protein were supplemented with 2nM FLAG-FANCD2. Pull-down was performed using FANCD2 antibody. Products were separated using SDS-PAGE and imaged by western blot analysis using 3XFLAG antibody. revealing both FANCD2, as indicated, and DNA2 full length and deletion proteins. (G) Mapping of the FANCD2 inhibitory domain-An N-terminal fragment F1 and C terminal fragment F4 of FANCD2 bind to DNA2. (H) The N terminal, DNA2-interacting domain of FANCD2 inhibits DNA2 nuclease. DNA2 nuclease assays were performed as in Figure 1 using the FANCD2 fragments indicated (Niraj et al., 2017). Assays were performed in duplicate using 0.2 nM DNA2 and 30 nM FANCD2 and were repeated four times. Ubi indicates the mutation of the ubiquitination site on F1.

To directly test this, we purified His-tagged hFANCD2 from *E. coli* and FLAG-tagged hDNA2 protein from human cells as described in Method Details (Takahashi et al., 2014) (Figure S2C). Immunoprecipitation experiments show that purified DNA2 and FANCD2 bind directly and strongly to each other (Figure 2C-D). Coimmunoprecipitation experiments show that the FANCD2/DNA2 interaction is independent of DNA (Figure 2E), further supporting that in vivo interaction may also be direct. These data are consistent with our previous finding that DNA2 and FANCD2 reciprocally co-IP in extracts of CPT-treated cells and that the interaction is independent of DNA (Karanja et al., 2012). Since the FANCD2 was prepared in *E. coli* for the in vitro experiments, it is not likely to be ubiquitylated, suggesting ubiquitin is not necessary for the interaction between FANCD2 and DNA2.

Using site-directed mutagenesis we identified a region on DNA2 in the N terminus spanning a.a. (amino acid) 227 to 348 that severely reduces coimmunoprecipitation with full-length FANCD2 (Figure 2F and Method Details). This region includes the canonical DEK nuclease family active site motifs (Budd and Campbell, 2000; Yang, 2011), suggesting how the interaction might interfere with nuclease function. Thus far, point mutations introduced into the active site region inactivate the catalytic activity of DNA2, so they have not been useful for further correlating the site in DNA2 required for nuclease inhibition by FANCD2.

In complementary experiments, we used a previously described complete set of contiguous fragments of FANCD2 (Niraj et al., 2017) to determine the region of FANCD2 that interacts with DNA2 and that inhibits DNA2, a functional assay for “interaction”. As shown in Figure 2G, fragment F1, a.a. 1-588, and fragment F4, a.a. 1178-1451, both coimmunoprecipitated with DNA2. This may identify an interface between these subunits that interacts with DNA2. However, fragment F1 was the only sub-fragment that inhibited DNA2 (Figure 2H). This region contains the FANCD2 ubiquitylation site (a.a. K561), and addition of a ubiquitin coding sequence to F1 (fragment designated F1-ubi, increased the efficiency of inhibition. We conclude that a protein/protein interaction is needed to inhibit DNA2 nuclease activity.

Since the FANCD2-F1+F3Mut protein showed residual binding to DNA, however, to further strengthen the conclusion that DNA2 inhibition is through a protein/protein interaction, we investigated whether inhibition by FANCD2 is species-specific. Yeast lacks a FANCD2 ortholog, and we hypothesized that yeast DNA2 would only be inhibited by FANCD2 if inhibition was mediated by occlusion/sequestration of DNA, thus preventing binding by DNA2. We observed no inhibition of yeast DNA2 by FANCD2, even at great molar excess FANCD2, on either the forked substrate or the reversed fork substrate (Figure S2D and S2E). Note that yDNA2 is more active than hDNA2, as also reported by others (Kumar et al., 2017), accounting for the concentrations used. The lack of inhibition of yeast DNA2 by FANCD2 supports, though it does not prove, that inhibition of hDNA2 by FANCD2 involves a species-specific and therefore likely a physiologically significant protein/protein interaction.

### DNA2 is also Inhibited by RAD51 Filaments

In addition to rescue by DNA2 nuclease depletion or specific chemical inhibition of nucleases, failure of BRCA2 or FANCD2 fork protection and resulting nascent DNA degradation has been shown to be rescued by elevated RAD51 levels or by stabilization of RAD51 filaments (Bhat et al., 2018; Schlacher et al., 2012; Wang et al., 2015). Furthermore, FANCD2 and RAD51 show epistatic interaction in nascent DNA degradation assays, i.e. destabilization of RAD51 filaments does not lead to further degradation of nascent DNA in FANCD2 deficient cells as determined by DNA fiber tracking (Schlacher et al., 2012) and see also(Hashimoto et al., 2010; Hashimoto et al., 2012; Higgs and Stewart, 2016; Kolinjivadi et al., 2017b; Schlacher et al., 2011; Schlacher et al., 2012; Taglialatela et al., 2017; Thangavel et al., 2015; Wang et al., 2015; Zadorozhny et al., 2017). To explore potential molecular interplay between FANCD2 and RAD51 in regulating DNA2-mediated resection, we first looked at whether there is physical interaction between DNA2, RAD51, and FANCD2 in HU-treated cells. We show that RAD51 co-IPs with FLAG-DNA2 and with endogenous FANCD2 (Figure 3A). Reciprocally, we immunoprecipitated RAD51 and showed that both FANCD2 and endogenous DNA2 coimmunoprecipitated (Figure S3A). The RAD51 immunoprecipitation in Figure S3A, which was analyzed under different gel conditions from the α-FLAG-DNA2 IP in Figure 3A, reveals that both mono-ubiquitylated and non-ubiquitylated FANCD2 are present in the co-immunoprecipitates. We then repeated these experiments after treating extract with Benzonase. The RAD51/FANCD2 interaction was still observed and is therefore not dependent on DNA (Figure 3B), in keeping with previous observations (Sato et al., 2016). However, since RAD51 was not found in a DNA2 IP after treatment of extracts with Benzonase (Figure 2E), we conclude the interaction of DNA2 with RAD51 is though DNA binding.

**Figure 3.**
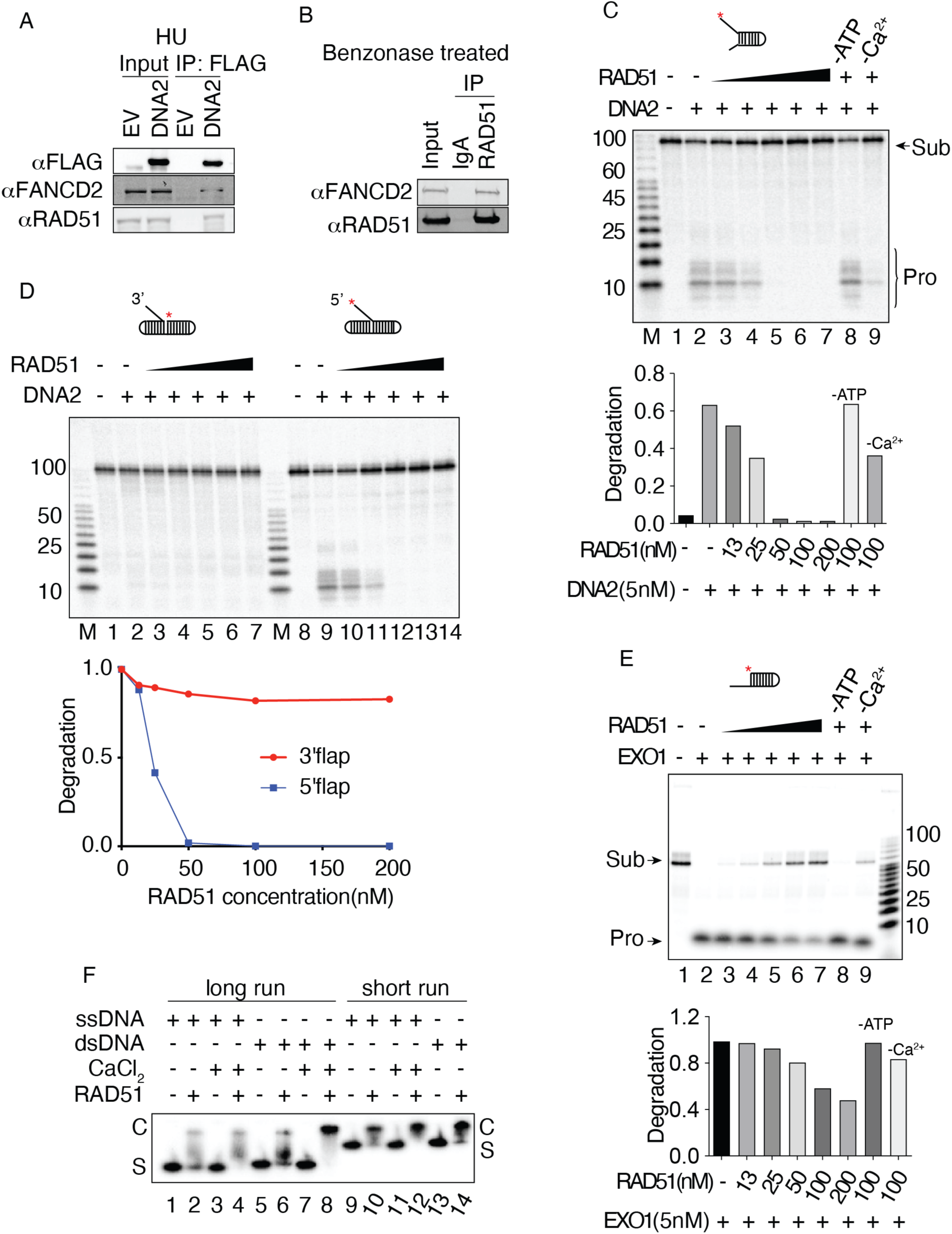
RAD51 filaments protect DNA from DNA2 mediated degradation. (A) Co-immunoprecipitation of FANCD2 and RAD51 with DNA2 (FLAG-DNA2 pull-down). FLAG-tagged DNA2 or empty vector was transfected into A549 cells, 24 hours later cells were treated with 2 mM HU for 3 hours and then harvested. Cells were lysed and pull down carried out with FLAG M2 beads; the beads were washed with lysis buffer then boiled with SDS loading buffer; immunoprecipitants were analyzed by western blotting of a 12% acrylamide gel with the indicated antibodies. (B) RAD51 and FANCD2 interact in the absence of DNA. U2OS cell nuclear extract was prepared as described in Method Details. Extracts were treated with Benzonase. 10 μl RAD51 antibody or IgG control added to 500μg nuclear extract and incubated for overnight at 4°C. 10ul Protein A agarose beads were added and then incubated for 1 hour at room temperature. The beads were washed with nuclease buffer and boiled with SDS sample buffer for western blot. (C) RAD51 filaments inhibit DNA2 on the forked substrate (87 FORK). Increasing amounts of RAD51 were preincubated with 4 nM 87 FORK substrate prior to the addition of 5nM DNA2. Controls show DNA2 activity in the presence of 100 nM RAD51 and in the absence of ATP (lane 8) or Ca^2+^ (lane 9). (D) RAD51 filaments inhibit DNA2 on a 5’ or 3’ flap. The indicated amounts of RAD51 were incubated with 4 nM of the respective flap substrate prior to the addition of 5nM DNA2. (E) RAD51 inhibits EXO1 nuclease on an overhang substrate. (F) Ca^2+^ enhances DNA binding activity of RAD51. 1 nM ssDNA or dsDNA was incubated in a 10 μl reaction mixture containing 200 nM RAD51, 25 mM TrisOAc (pH 7.5), 1 mM MgCl_2_, 2 mM CaCl_2_ and 2 mM ATP (except where CaCl_2_ was omitted), 1 mM DTT, 0.1 BSA mg/ml, as indicated. The reaction was incubated for 10 min at 37°C(Wang et al., 2015) and samples were mixed with 1 μl loading buffer (2.5%Ficoll-400, 10 mM Tris-HCl, pH 7.5 and 0.0025% xylene cyanol). Products were analyzed on a 5% native gel (29:1 30% acrylamide in TAE), constant voltage, 60V, in the cold room for 1h (lanes 9-14) or 2h (lanes 1-8) followed by phosphor imaging.

Although DNA2 and RAD51 don’t appear to interact stably in the absence of DNA, RAD51 filaments have been implicated in regulating DNA2-mediated resection, since RAD51 T131P, which fails to form stable filaments, leads to over-resection in heterozygous FANCR patients bearing the mutation and the over-resection is reversed by DNA2 depletion (Wang et al., 2015). We tested for inhibition of FLAG-DNA2 nuclease by recombinant RAD51 protein. Increasing amounts of RAD51 inhibited DNA2 nuclease activity on both fork and flap substrates (Figures 3C and 3D). RAD51 also inhibits EXO1 nuclease on an overhang substrate (Figure 3E) and has previously been shown to inhibit MRE11 (Kolinjivadi et al., 2017b). This general inhibition might be attributed to inhibition by filament formation blocking the nucleases. To distinguish between inhibition by RAD51 protein alone and inhibition by RAD51 filament formation, we took advantage of the demonstration that ATP but not ATP hydrolysis is required for filament formation and that Ca^2+^ permits filament formation but modulates ATP hydrolysis and inhibits dissolution of the filaments, resulting in stabilization (Bugreev and Mazin, 2004). As shown in Figure 3C and 3E, RAD51 does not inhibit nucleases in the absence of ATP and Ca^2+^ stimulates inhibition (Figures 3C, 3E-F, Figure S3B). As verified in Figure 3F and S3B, RAD51 filaments were formed on both ssDNA and dsDNA in the presence of ATP and are more stable in the presence of Ca^2+^ than in its absence. We conclude that inhibition of DNA2 is mediated by RAD51 filaments.

### FANCD2 Stimulates Strand Exchange by High Concentrations of RAD51

We were struck by the fact that BRCA2^−/−^ cells and FANCD2^−/−^ show non-epistatic interactions such as synthetic lethality and that over-expression of FANCD2 suppresses BRCA^−/−^ phenotypes (Kais et al., 2016; Michl et al., 2016). Furthermore, like BRCA2, FANCD2 interacts physically and robustly with RAD51 [Figure 3 and (Chen et al., 2017; Sato et al., 2016)], and RAD51 has been shown to localize to stalled forks in cells lacking BRCA2 (Kolinjivadi et al., 2017b). We hypothesized that FANCD2 might, similarly to BRCA2, stimulate RAD51-mediated strand exchange (Jensen et al., 2010; Thorslund et al., 2010). While FANCD2 does not enhance RAD51-mediated D-loop assays with resected plasmid substrates (Dubois et al., 2019), complete strand exchange assays with oligonucleotides were never tested. Both reactions are linked to DNA recombination, but mechanistically they are different. In D-loop assays strand invasion into a supercoiled DNA recipient is measured and is thought to represent a search for homology (Carreira et al., 2009; Carreira and Kowalczykowski, 2011). Strand exchange assays, in contrast, measure a complete transfer of DNA strands (see schematic in Figure 4A). As indicated, RAD51 catalyzes the exchange of the labeled strand in the duplex to ssDNA to form the strand exchange product (Jensen et al., 2010; Thorslund et al., 2010). High concentrations of RAD51, however, have been shown to be inhibitory in this assay (Jensen et al., 2010; Thorslund et al., 2010). To measure strand exchange, RAD51 was incubated, in the presence or absence of FANCD2, with ssDNA (Figure 4B, pilot experiment), with duplex DNA with a 3’ ssDNA overhang (Figure 4C), or with duplex DNA with a 5’ ssDNA overhang (Figure 4C) to allow filament formation. Fully duplex DNA containing a ^32^P labeled strand complementary to the ssDNA or respective overhang DNA was then added. Stimulation of strand exchange by RAD51 is shown for ssDNA in Figure 4B, lanes 1-4. Inhibition at high RAD51 levels is shown in Figure 4B, lane 5. Such inhibition is proposed to arise once ssDNA is saturated with RAD51, allowing the excess RAD51 to bind to the labeled dsDNA donor, which inhibits exchange (Jensen et al., 2010; Thorslund et al., 2010). Supporting the hypothesis that the inhibition by high levels of RAD51 can be due to binding of excess RAD51 to duplex DNA, we showed that addition of a dI-dC oligonucleotide relieves inhibition, presumably by successfully competing with the labeled duplex donor for excess RAD51 binding in the assays (Figure 4B, lane 12). We then studied whether FANCD2 stimulated RAD51 at high RAD51 concentrations, as has been shown for BRCA2 (Jensen et al., 2010; Thorslund et al., 2010). As shown in Figure 4B (lanes 6-11) and Figure 4C, although FANCD2 has no strand exchange activity on its own, FANCD2, indeed, reproducibly stimulates strand exchange by high concentrations of RAD51 and does so in a concentration dependent manner. Strand exchange involving duplex DNA with a 3’ or 5’ overhang, more closely resembling a filament on resected DNA, was stimulated more efficiently than with ssDNA, suggesting that stimulation may occur on DNA with ds/ss junctions and may occur at gaps as well as at ssDNA tails (Figure 4B, C). Several controls that strand exchange was occurring were performed. Reversing the order of addition of substrates, i.e., formation of RAD51 filaments on dsDNA and then addition of ssDNA, did not lead to exchange (Figure 4D); thus, we are not observing inverse strand exchange (Mazina et al., 2017). Addition of cold oligonucleotide to the stop reaction does not change the products, supporting that the strand exchange products are not formed due to denaturation and renaturation in the stop mixture (Figure 4C, lanes labeled cold oligo)(Jensen et al., 2010). We conclude that FANCD2 stimulates RAD51 mediated strand exchange.

**Figure 4.**
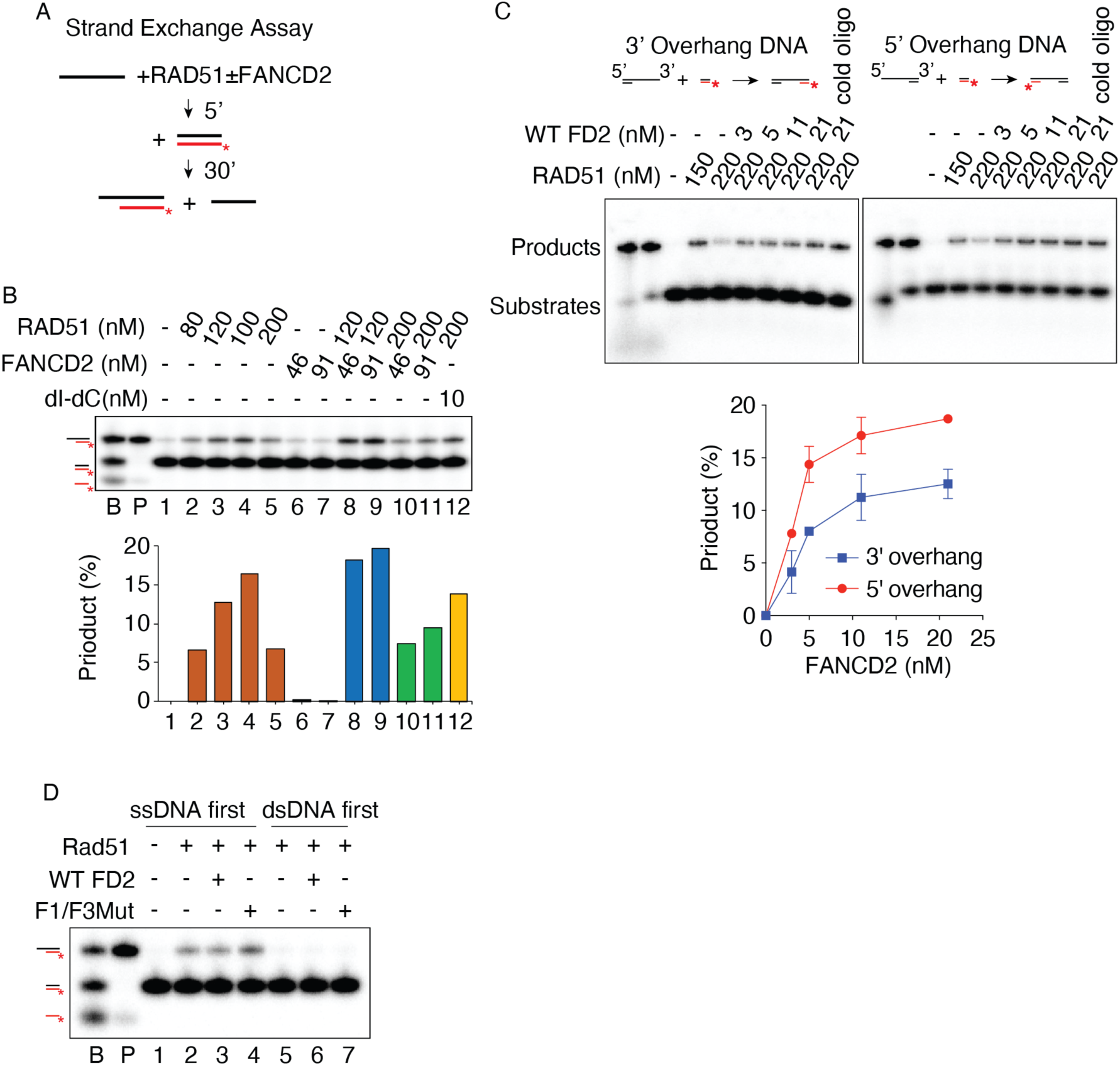
FANCD2 stimulates RAD51-mediated strand exchange. (A) Schematic of strand exchange assay: Single-stranded or 3’ or 5’ overhang DNA is incubated in the presence of RAD51 to form filaments. The filaments are then incubated with a duplex DNA with a labeled strand complementary to the filament. Product formation, representing complete strand exchange, is monitored using a native acrylamide gel. (B) FANCD2 stimulates strand exchange on ssDNA by high levels of RAD51. Quantification is shown below each gel. Lane labeled B shows the migration of each relevant DNA species as indicated in the schematic on the left and was prepared by annealing oligonucleotide EXTJYM925, JYM925, and 5’ labeled JYM945; the lane labeled P confirms the position of the exchanged strand product (EXTJYM925 and 5’ labeled JYM945). Lanes 1-5: 4 nM ssDNA (100 nt, oligonucleotide EXTJYM925) was incubated with indicated amounts of RAD51 for 5 min at 37°C and the 5’ labeled dsDNA (60mer, JYM925/JYM945 oligonucleotides) (final concentration 4 nM) was added and incubation continued for an additional 30 min at 37°C for strand exchange. Lanes 6 and 7, as in lanes 1-5 with indicated concentrations of FANCD2 in the absence of RAD51. Lanes 8-11: RAD51 plus FANCD2 at the indicated concentrations present during both the 5’ preincubation with ssDNA and after addition of dsDNA. Lane 12: 10 nM of dI-dC competitor present during preincubation of RAD51 and ssDNA. Histogram shows quantitation. (C) FANCD2 stimulates strand exchange on 3’ and 5’ overhang DNA by high levels of RAD51. Reactions performed as in panel B; however, 4 nM 3’ overhang DNA (162 nt RJ-167 annealed to 42 nt RJ-PHIX-42-1) (Left) or 4 nM 5’ overhang DNA (162 nt RJ-167 annealed to 42 nt RJ-PHIX-42-2) (right) as indicated, were incubated in the presence of indicated amounts of RAD51 for 5 min at 37°C to form filaments. 5’ labeled dsDNA (40mer, RJ-Oligo1/RJ-Oligo2 in the case of 3’ overhang DNA or RJ-Oligo4/RJ-Oligo3 in the case of 5’ overhang DNA) (final concentration 4 nM) was added and incubation continued for an additional 30 min at 37°C for strand exchange. Lane 1, no protein; lanes 2-3: RAD51 alone; lanes 4-7: RAD51 plus FANCD2 at the indicated concentrations present during both the 5’ preincubation with 3’ or 5’ overhang DNA and after addition of dsDNA. Lane 8: 10-fold excess of unlabeled heterologous ssDNA (40 nt, oligonucleotide RJ-Oligo2) complementary to labeled strand of dsDNA was added to the stop solution to rule out that the product observed was due to denaturation and annealing during the deproteinization/termination step. (% Product represents the value with unstimulated exchange subtracted.) The graph shows quantification for both assays. The assays were repeated twice. (D) Inverse strand exchange assay. In lanes 1-4, the exchange assay was conducted as in the legend to Figure 4A-C. In lanes 5-7 the double-stranded DNA was preincubated with RAD51 and then ssDNA was added.

### FANCD2 Promotes Strand Exchange Activity by Enhancing ssDNA Binding of RAD51

We next interrogated the mechanism of FANCD2 stimulation of RAD51. BRCA2 DNA binding is required for stimulation of strand exchange, and BRCA2 is thought to stimulate strand exchange in several ways: by stabilizing RAD51 filaments through inhibiting RAD51 DNA-dependent ATPase, by promoting the handoff of ssDNA from RPA to RAD51, and by nucleating filament formation on ssDNA while inhibiting filament formation on duplex DNA. To determine if FANCD2 DNA binding was required for stimulation of strand exchange, we tested if the FANCD2 DNA binding mutant described above stimulated strand exchange (Niraj et al., 2017). Although FANCD2-F1+F3Mut showed approximately ten-fold reduction in ssDNA binding at 10 nM (Figure S2B), FANCD2-F1+F3Mut protein stimulates strand exchange even more efficiently than FANCD2 WT (Figure 5A), which suggests DNA binding activity of FANCD2 is not required in promoting strand exchange, or that weak binding is sufficient. We next determined if FANCD2 inhibits RAD51 DNA-dependent ATPase. Surprisingly, unlike BRCA2, FANCD2 does not inhibit RAD51 DNA-dependent ATPase (Figure 5B), and thus may not be acting to stabilize RAD51/ssDNA filaments by blocking the ATPase.

**Figure 5.**
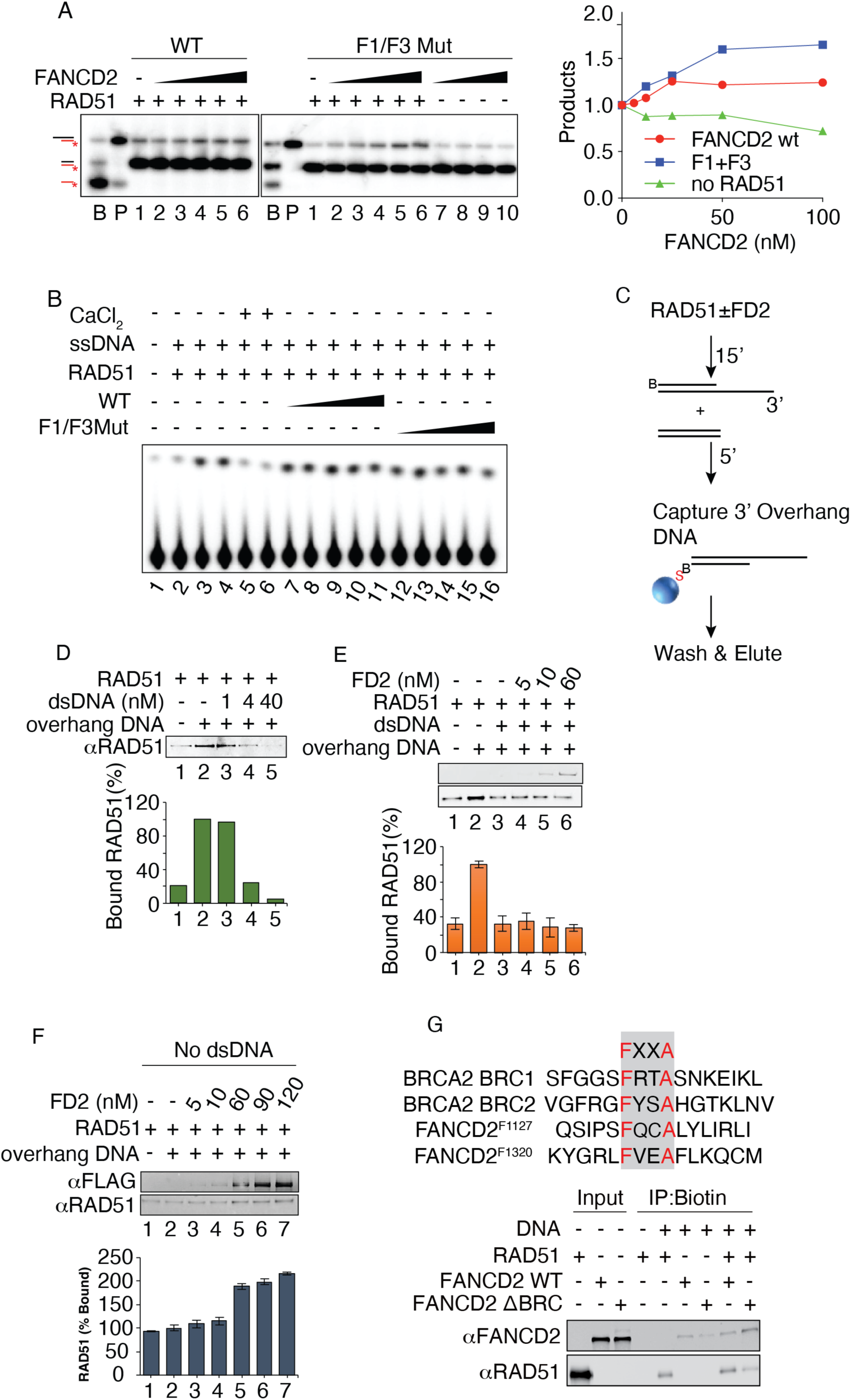
FANCD2 stimulates strand exchange activity by enhancing ssDNA binding of RAD51. (A) Both wild-type FANCD2 and FANCD2-F1+F3M stimulate strand exchange. Lanes 1-6 (left): Titration of WT FANCD2 at 100 nM RAD51 and 2 nM of both ssDNA and dsDNA. Assays were performed with oligonucleotides as in panel B. Lanes 2-10 (right): Indicated amounts of FANCD2-F1+F3 mutant incubated in the absence of RAD51 during both the preincubation with ssDNA and after addition of dsDNA. Graph shows quantification for both assays. (B). Wild type FANCD2 and FANCD2-F1+F3 mutant do not inhibit the DNA-dependent ATPase activity of RAD51. FANCD2 WT and F1+F3 mutant concentration is 6, 11, 23, 46, 91 nM. 300 nM RAD51 and 900 nM ssDNA added to the reaction. (C) Schematic of biotinylated DNA pull-down assay, B: B-biotin; S: S-streptavidin. (D) Assembly of RAD51 onto 3’ overhang DNA is suppressed by heterologous dsDNA competitor. RAD51 and FANCD2 proteins at indicated concentrations were incubated for 15 min at 37°C followed by the addition of 3’ overhang DNA (162 nt RJ-167 annealed to 42 nt 3’ Bio-RJ-PHIX-42-1) and competitor heterologous dsDNA (90mer, Oligo #90/Oligo #60 oligonucleotides) and incubated for an additional 5 min at 37°C. Where DNA was omitted, TE buffer was used and similarly, respective proteins storage buffers were used where proteins were omitted. 20 µL reactions were performed as in panel f using 60 nM RAD51 either in the absence (lane 2) or presence (lanes 3-5) of excess competitor heterologous dsDNA (90mer, Oligo#90/Oligo#60 oligonucleotides). Histogram shows quantification. Assays were repeated two times. (E) FANCD2 does not rescue RAD51 filament formation on 3’ overhang DNA in the presence of heterologous dsDNA competitor. 20 µL reactions were performed as in panel D using 60 nM RAD51 preincubated with (lanes 3-6) or without (lane 2) increasing concentrations of FANCD2 (FD2). Both proteins were separately probed by western analysis. The histogram shows quantification of the RAD51 western analysis. Assays were repeated two times. (F) FANCD2 stimulates RAD51 filament formation on 3’ overhang DNA in the absence of dsDNA competitor. Reactions were performed as in panel E using 60 nM RAD51 preincubated with (lanes 3-7) or without (lane 2) increasing concentrations of FANCD2. TE Buffer in lieu of dsDNA was added with 3’ overhang DNA for all samples and incubated for an additional 5 min at 37°C. Both proteins were separately probed for western blot analysis. Histogram shows quantification of the RAD51 western blot analysis. Assays were repeated two times. (G). FANCD2 stimulates recruitment of RAD51 to DNA but BRC repeat double mutant of FANCD2 inhibits recruitment of RAD51 to DNA. Top: map positions of FXXA motifs in BRCA2 and FANCD2. Bottom: The conditions were the same as for panel F.

We finally tested if FANCD2 plays a role in targeting RAD51 preferentially to ssDNA by inhibiting nucleation on dsDNA. We carried out DNA binding experiments using biotin-streptavidin pull-downs (Figure 5C). We first demonstrated that dsDNA inhibits RAD51 binding to a biotin-labeled 3’ overhang substrate (Figure 5D). We then added FANCD2 and found that FANCD2 did not stimulate association of RAD51 with the 3’ overhang substrate in the presence of excess dsDNA (Figure 5E). This is unlike what has been demonstrated for BRCA2, which has been shown to specifically overcome dsDNA inhibition of RAD51 binding to overhang DNA in a similar assay (Jensen et al., 2010; Thorslund et al., 2010), Thus, FANCD2 is not stimulating RAD51 by reducing binding to dsDNA and is more likely stabilizing RAD51/ssDNA filaments. Supporting this interpretation, FANCD2 alone does stimulate the accumulation of RAD51/ssDNA complexes in the absence of dsDNA (Figure 5F), similar to previous characterization of RAD51/FANCD2/FANCI interaction with DNA (Sato et al., 2016). BRC repeats have been shown to be required for stimulation of strand exchange by BRCA2. The FANCD2 protein carries two FXXA BRC consensus site motifs, at F1127 and F1320, respectively (Carreira et al., 2009; Carreira and Kowalczykowski, 2011; Jensen et al., 2010). To test if they were required to stimulate RAD51 DNA binding, a FANCD2 F1127A/F1320A mutant DNA was constructed and the mutant protein expressed and purified. As shown in Figure 5G, the wild-type FANCD2 protein stimulated RAD51 DNA binding in the biotin pull-down assay but the BRC-mutant FANCD2 protein reproducibly inhibited binding of RAD51. Based on the results in Figures 4 and 5, we suggest that FANCD2 stimulates strand exchange by directly promoting, either through nucleation, assembly, or filament stabilization, RAD51/ssDNA filament formation, rather than by competing with dsDNA. This FANCD2-mediated stabilization does not involve inhibition of RAD51 ATPase and does not require optimum FANCD2 DNA binding activity. Stimulation of strand exchange with these characteristics suggests that the molecular role of FANCD2 in fork protection may involve stimulation of formation of or stabilization of RAD51 filaments, and thus indirect inhibition of DNA2 nuclease, in addition to the direct inhibition of DNA2 shown in Figure 2. The results further suggest that interaction of FANCD2 with RAD51 protein [Figure 3 and (Sato et al., 2016)] contributes to strand exchange, and therefore that this may contribute to the ability of elevated levels of FANCD2 to suppress some BRCA2-deficiencies in fork protection (Ceccaldi et al., 2015; Michl et al., 2016).

### FANCD2 May Play Different Roles at Different Types of DNA Damage

As shown in Figure 1A, we observed substantial over-resection in the absence of FANCD2 after HU treatment of cells. This suggests that the damage, likely consisting of helicase/polymerase uncoupling and fork reversal damage subsequent to stalling (Byun et al., 2005; Taylor and Yeeles, 2019; Zellweger et al., 2015), is protected from resection by FANCD2. Such a role for FANCD2 in negative regulation of resection, however, needs to be reconciled with previous elegant studies showing that FANCD2 is actually required for resection for repair after extensive ICL-induced stalling, which may induce a different type of damage, including DSBs and or gaps (Murina et al., 2014; Unno et al., 2014; Yeo et al., 2014).

To support the proposal that FANCD2 can have two opposing effects on resection in response to different types of damage, we carried out time courses of CPT (camptothecin) treatment in PD20 or FANCD2 depleted cells and respective FANCD2-complemented cells. While low levels of CPT can simply induce fork slowing or stalling (through topological stress) (Ray Chaudhuri et al., 2012), high dose CPT rapidly induces DSBs when the replication fork encounters sites of the CPT-induced Top1-DNA cleavage complexes (Berti et al., 2020; Whelan and Rothenberg, 2021). We observed that in the absence of FANCD2, there is over-resection at the earliest time point (Figure 6A, lanes 3 and 4 compared to lanes 5 and 6 and Figure 6B). At later times, FANCD2 becomes a positive regulator of resection, however, presumably at rapidly accumulating “collapsed forks”, since there is reduced resection in the absence of FANCD2 (Figure 6A-C). Cells lacking FANCD2 respond to extensive cisplatin treatment similarly to CPT (Figure 6C). Neutral COMET assays support a greater abundance of DSBs in CPT treatment than in HU and a more rapid increase in DSBs during CPT treatment than in HU (Figure 6D-E). We cannot distinguish whether the initial damage is being remodeled during the time course or if different structures arise independently as stalled forks are remodeled in response to damage. Taken together, our results suggest that FANCD2 is required to protect from over-resection after damage on forks transiently stalled by HU, probably largely on reversed forks or gaps. FANCD2, however, may play a different role during chronic stalling that leads to fork collapse to DSBs or to other types of damage, such as gaps due to Prim-Pol activity, when FANCD2 may actually be required for resection and repair (Figure 6A-B). This interpretation is consistent with recent proposals that different repair mechanisms may function on HU and CPT damage (Couch et al., 2013; Liu et al., 2020; Rickman et al., 2020).

**Figure 6.**
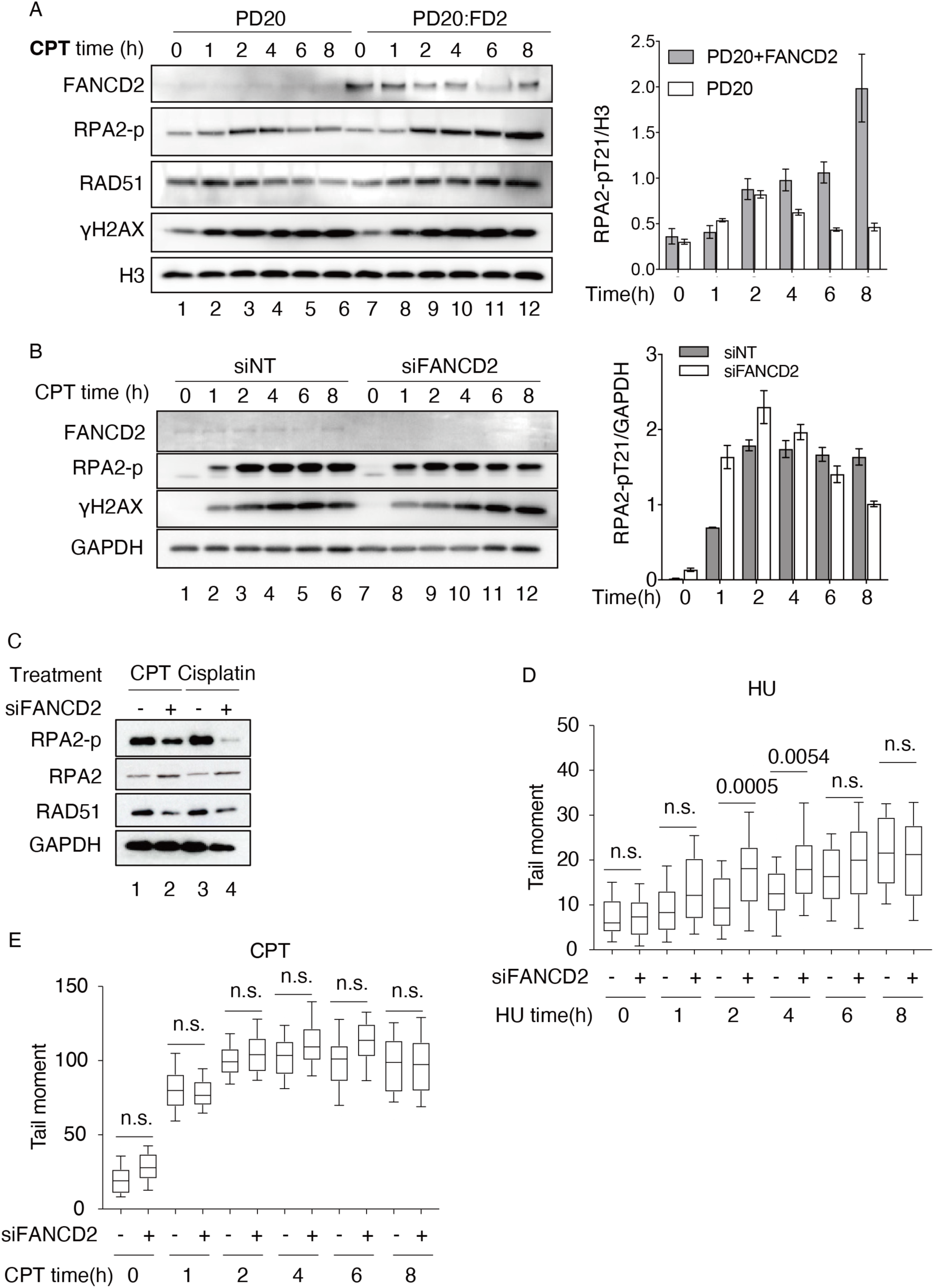
FANCD2 acts differently on different types of DNA damage. (A) FANCD2 is required for resection in CPT treated cells as revealed by time course of treatment of PD20 FANCD2-/- cells with CPT. Assays for resection are as in Figure 1A. PD 20 FANCD2^−/−^ cell and PD20 complemented with FANCD2 (PD20:FD2) were treated with 2 μM CPT for indicated times. Nuclear extract was prepared and used for the western blots for proteins and protein modifications as indicated in the figure. (B) FANCD2 is confirmed to be required for resection of CPT treated cells as revealed by knockdown of FANCD2. A549 cells were transfected with 40 nM FANCD2 siRNA or control scrambled siRNA. After 48 h, 2 μM CPT was added and cells were treated for 0-8 hours. Samples were taken and whole cell lysates (GAPDH control) were prepared for western blot analysis after 0, 2, 4, 6, or 8 h of treatment. RPA2-p and γH2AX were monitored as shown. The RPA2-p bands were quantified with ImageJ, and error bars derive from standard deviation of two experiments. Resection was measured as RPA-p levels on western blots, as in Figure 1. (C) FANCD2 is required for resection in cisplatin treated cells after prolonged exposure.A549 cells were transfected with 40 nM FANCD2 siRNA or control scrambled siRNA (siNC). After 48 h, 2 μM CPT or 10 mM cisplatin was added as indicated and cells incubated for 16h. Samples were taken and whole cell lysates (GAPDH control) were prepared for western blot. RPA2-p and γH2AX were monitored as shown. (D and E). Neutral comet assay of DNA damage in HU treated cells compared to CPT-treated cells over time suggests greater number of DSBs in the CPT treated cells.

## Discussion

FANCD2 has been studied for decades and much has been learned about its cellular functions and structure. What is unknown, however, despite extensive cellular and structural characterization, are the biochemical activities it uses to accomplish and coordinate its diverse in vivo roles. We and others have shown that FANCD2 is a fork protection factor that prevents MRE11 and DNA2 mediated resection at replication forks upon HU fork stalling and remodeling into reversed forks. The replication fork protection function of FANCD2 may be distinct from its canonical roles in the Fanconi anemia pathway since different substrates may be involved. How FANCD2 protects the stalled fork and what kind of DNA structures it acts upon remain unclear. Here, our study utilizes biochemical assays to uncover three ways that FANCD2 protects the stalled forks or gaps: 1) FANCD2 directly binds to DNA2’s nuclease domain to inhibit its nuclease activity; 2) FANCD2 stabilizes RAD51 filaments to prevent non-specific degradation by multiple nucleases; 3) Apart from regulating nascent strand degradation, FANCD2 stabilizes RAD51 filaments on ssDNA to stimulate strand exchange activity, which may parallel BRCA2, explaining the previous finding that FANCD2 compensates for BRCA2’s loss.

### FANCD2 Directly Inhibits DNA2 in Vitro Identifying a Non-canonical Role for FANCD2 in Protecting Stalled Forks from Degradation

Our in vitro nuclease inhibition assay may indicate that FANCD2 prevents over-resection at stalled forks independent of its roles in ICL repair. Several observations are consistent with the conclusion: 1) FANCI did not efficiently stimulate the FANCD2-mediated inhibition of DNA2, nor did FANCI inhibit DNA2 significantly on its own. Failure to stimulate FANCD2 inhibition might be explained if the FANCD2 that inhibits DNA2 is in a dimeric form, which has also been reported to fail to interact with FANCI (Alcon et al., 2020). Our finding does not exclude the possibility, though, that FANCD2/I heterodimers may participate when present. 2) FANCD2 stably and specifically binds DNA2. Ubiquitylation is not essential for interaction but seems to augment inhibition (Figure 2). 3) FANCD2 doesn’t suppress the nuclease activity of the heterologous yeast DNA2, further suggesting that FANCD2/DNA2 protein/protein interaction is important in downregulating the nuclease. We identified two DNA2 interaction domains in FANCD2, the F1 and F4 fragments of FANCD2. Furthermore, we found that FANCD2-F1, but not FANCD2-F4, inhibits DNA2. In a complementary experiment, we showed that the FANCD2-interaction domain within DNA2 lies in the N-terminal region comprising its nuclease catalytic site. Thus, we propose that the direct binding of FANCD2-F1 to the nuclease domain of DNA2 may hinder the nuclease activity. 4) FANCD2 inhibits DNA2 even when FANCD2 DNA binding is compromised. Recent reports found that FANCD2 is purified as a dimer (Tan et al., 2020) and suggested that the dimer is not capable of DNA binding (Alcon et al., 2020; Alcon et al., 2019). However, our wild-type FANCD2 preparation does bind dsDNA (Figure S1B). It was proposed that the DNA binding defect in the FANCD2 dimers might be due to sequestration of the DNA binding domain (Alcon et al., 2020; Alcon et al., 2019). We do not know if our preparation contains monomeric or dimeric FANCD2, as gel filtration experiments were inconclusive to date.

The role of FANCD2 in inhibiting DNA2 over-resection may be important for replication restart. Replication forks are constantly challenged by various stresses, causing them, if not rapidly repaired, to be converted to damaged forms in need of nucleolytic processing. Such processing, however, is a dangerous game. It must be highly regulated to prevent aberrant processing giving rise to DNA breaks, chromosomal rearrangements, and aneuploidy, which are hallmarks and drivers of cancer. Paradoxically, numerous anti-cancer agents kill cancer cells by impairing the regulated progression of the replication machinery and inducing replication stress. On the other hand, controlled resection, in other words, fork protection, may lead to chemoresistance. Thus, cells have evolved multiple pathways to modulate the nucleolytic processing and a host of protection factors have been identified – RECQ1, RAD51, BRCA1/2 and related factors along with multiple nucleases. Understanding the mechanism of fork protection in each case will help us to address drug resistance. As we have shown here, inhibition by FANCD2 represents a new aspect of the regulation of DNA2, which has already been shown to be modulated by multiple other mechanisms [see(Zheng et al., 2020) for a recent review].

DNA2 inhibitors (DNA2i) synergize with PARP inhibitors (PARPi) in killing BRCA-deficient breast cancer cells (Liu et al., 2016). Since DNA2 is required for DSB repair, the enhancement of PARPi sensitivity by DNA2i has been ascribed to the putative selectivity of PARPi for the DSB repair phenotype of BRCA-deficient cells (Cong et al., 2021). However, PARP is also implicated in protecting from ssDNA damage (Cong et al., 2021; Hanzlikova et al., 2018; Vaitsiankova et al., 2022). Thus, over-resection at ssDNA damage may also contribute to DNA2i and PARPi synergy. Specifically, on the lagging strand, PARP regulates a back-up mechanism for defective Okazaki fragment processing that can lead to an accumulation of unreplicated gaps (Hanzlikova et al., 2018; Vaitsiankova et al., 2022), possibly involving downstream pol α-primase repriming (Kolinjivadi et al., 2017b). PARP has been demonstrated to be involved in protection from excessive DNA2-dependent degradation of nascent DNA on the lagging strand (Thakar et al., 2019; Thakar et al., 2020), implying a need for DNA2 in processing lagging strand restart intermediates. ssDNA gaps also arise on the leading strand, due to repriming by PRIMPOL (Quinet et al., 2019). Thus over-resection by DNA2 in the absence of FANCD2 may be deleterious not only during fork reversal and DSB repair but also at lagging or leading strand gaps. In keeping with this proposal, DNA2 shRNA knockdown leads to lengthened replication tracts in DNA fiber studies (Duxin et al., 2012). Such tracts could be due to aberrant gapped daughter chromosomes (Cong et al., 2021). Both PARPi and extensive treatment with cisplatin in a FANCD2 knockdown (Lossaint et al., 2013; Yeo et al., 2014) cause apparent fork speed acceleration, and enhanced replication fork speed can in itself cause genome instability and may involve unligated Okazaki fragments (Cong and Cantor, 2022; Maya-Mendoza et al., 2018; Merchut-Maya et al., 2019). Our results suggest a role for FANCD2 in modulating DNA2 resection in all of these stalled replication fork intermediates.

Recently, FANCD2 inhibition of DNA2 resection, surprisingly, has been demonstrated to play a significant role in restraining accumulation of cytoplasmic DNA and induction of the cGAS-STING pathway in response to replication fork stalling in vivo (Emam et al., 2022). Possible roles for FANCD2 in immune secretion and inflammation and their emerging link to DNA repair pathways lend further significance to understanding the mechanism by which FANCD2 controls DNA2.

### FANCD2 also Regulates Resection Through RAD51

Previously, the FANCD2/I heterodimer was shown to stabilize RAD51 filaments, but we have demonstrated that FANCD2 alone can stabilize RAD51 on ssDNA and that ablation of FXXA motifs found in putative BRC motifs within FANCD2 prevents RAD51 filament stabilization (Figure 5). Recent studies of replication initiation have revealed that even low levels of stabilization can have critical regulatory outcomes in the cell (Stillman, 2022). This uncovers a mechanism by which RAD51 overexpression, in addition to FANCD2, may suppress the DNA2-dependent component of nascent DNA degradation in BRCA2 and FANCD2 deficient cells (Schlacher et al., 2011; Schlacher et al., 2012). Like FANCD2, we showed that RAD51 filaments also inhibit DNA2 nuclease, and this requires stable RAD51/ssDNA filaments (Figure 3). We suggest that RAD51 filaments may more generally retard degradation by multiple nucleases, since we found that RAD51 filaments also protect from EXO1-mediated nascent DNA degradation, and others have demonstrated in vitro inhibition of MRE11 nuclease by RAD51 (Hashimoto et al., 2010; Kolinjivadi et al., 2017a; Kolinjivadi et al., 2017b).

We reasoned that the nuclease inhibition by RAD51 filaments was not the sole role of FANCD2 at stalled forks, given complex roles that RAD51 plays, including fork reversal. The fact that FANCD2 and BRCA2 are synthetically lethal and that FANCD2 overexpression suppresses the replication fork protection defect of BRCA^−/−^ cells, suggested that FANCD2 might have a parallel function to BRCA2 (Kais et al., 2016; Michl et al., 2016). Therefore, we tested whether FANCD2 had activities similar to BRCA2 protein, a known RAD51 mediator. We found that FANCD2 stimulates RAD51 strand exchange, overcoming inhibition of strand exchange by high levels of RAD51 and does so with similar stoichiometry to that reported for BRCA2 (Jensen et al., 2010; Thorslund et al., 2010). BRCA2 stimulates strand exchange on one level by acting as a mediator in the exchange of RPA for RAD51 on resected overhangs. Second, BRCA2 promotes RAD51 ssDNA filament formation through inhibition of non-productive or inhibitory binding of RAD51 to dsDNA, presumably by competition between BRCA2 and RAD51 for DNA binding (Jensen et al., 2010; Thorslund et al., 2010). BRCA2 uses BRC repeats 1-4 to inhibit RAD51 ATPase and thus to stabilize filaments, but BRCA2 also uses BRC repeat 6-8 to promote nucleation of RAD51 on ssDNA and thus stimulate strand exchange (Carreira et al., 2009; Carreira and Kowalczykowski, 2011). We did not find that FANCD2 inhibited RAD51 ATPase, nor did it overcome the inhibition of RAD51 binding to ssDNA by dsDNA (biotin pull-down assays). Thus, FANCD2 is acting differently from BRCA2 or MMS22L/TONSL (Piwko et al., 2016), which also stimulates strand exchange. Furthermore, the DNA-binding-defective FANCD2-F1+F3Mut protein, was even more efficient than WT FANCD2 in enhancing strand exchange, also differing from BRCA2, which needs to bind to DNA to stimulate RAD51. The (at least partial) independence of FANCD2 DNA binding in strand exchange suggests that stimulation of strand exchange by FANCD2 involves a significant FANCD2/RAD51 protein/protein interaction and that this in turn helps stabilize RAD51 filaments. Interestingly, the FANCD2/FANCI complex stabilizes RAD51 filaments, and FANCI DNA binding motifs are necessary but the FANCD2 DNA binding motifs are not necessary (Sato et al., 2016), in accordance with our observation that FANCD2 DNA binding mutants stimulate strand exchange even in the absence of FANCI. Stimulation of strand exchange by FANCD2 most likely involves stabilization in some way of the RAD51 filament, perhaps by preventing end release, as suggested previously for the FANCD2/FANCI complex (Sato et al., 2016) or by altering the filament structure in multiple ways, as demonstrated for RAD51 paralogs (Belan et al., 2021; Berti et al., 2020; Cejka, 2021; Liu et al., 2011; Roy et al., 2021; Sullivan and Bernstein, 2018; Taylor, 1958; Taylor et al., 2015). This proposal is supported by the fact that FANCD2 carrying mutations affecting two FXXA consensus BRC motifs, such as are found in the BRC1 and BRC2 motifs of BRCA2, fail to stimulate RAD51 binding.

One likely mechanism of FANCD2 stimulation of RAD51 filament formation is to provide a chaperone for RAD51 filament assembly. FANCD2, namely, has been shown to have a histone chaperone function in nucleosome assembly. The histone chaperone function of FANCD2 is stimulated by histone H3K4 methylation mediated by BOD1L and SETD1A. Strikingly, in BOD1L or SETD1A depleted cells or in cells with inactivated FANCD2 chaperone function, RAD51 filaments are destabilized and stalled forks are excessively degraded by DNA2 defective (Higgs et al., 2018). Our results are consistent with the suggestion that FANCD2 might assist stable complex formation between RAD51 and DNA, i.e. filament nucleation, elongation or stabilization, in addition to promoting histone association and appropriate chromatin structure to protect stalled forks. A similar chaperone-like function has also been proposed for the RAD51 paralogs (Belan et al., 2021; Cejka, 2021; Halder et al., 2022; Roy et al., 2021), and many histone chaperones have been shown to chaperone additional proteins into assemblies,

### Possible Roles of FANCD2-stimulated Strand Exchange Activity in Fork Protection

How does strand exchange support fork protection and restart of forks? There are several possibilities (Bhat and Cortez, 2018). 1) Stabilizing RAD51 filaments on the ssDNA arising at uncoupled replication forks could stimulate strand exchange promoting fork reversal processes (Berti et al., 2020; Mijic et al., 2017; Piwko et al., 2016; Zellweger et al., 2015), 2) Another possibility is that RAD51 could promote reannealing of the ssDNA immediately behind the fork, essentially zipping up the unwound DNA, helping to promote fork reversal. Fork reversal could slow replication forks for repair; 3) the reversed forks might be the substrates for FANCD2-controlled DNA2-mediated processing, leading to replication fork restart (Thangavel et al., 2015). Two studies did show that cells deficient in FANCD2 failed to restrain synthesis in the presence of HU or aphidicolin, possibly by failing in fork reversal (Lossaint et al., 2013; Yeo et al., 2014). 4) FANCD2 stimulation of RAD51 strand exchange may be important for post-replication repair (Petermann et al., 2010; Piberger et al., 2019). While it was originally thought that the structure of the forks at these extensively damaged chromosomes might be DSBs, recent evidence suggests that they may also be unrepaired gaps after repriming, by Prim-Pol, at leading strand blocks or by pol α–primase at blocks on the lagging strand, especially (Cong and Cantor, 2022; Quinet et al., 2019). FANCD2 might stimulate RAD51-mediated strand exchange events in post-replication repair and template switch at these sites, especially if BRCA2 is defective (Fumasoni et al., 2015). 5) In BRCA2^−/−^ cells excessive resection creates a substrate that requires MUS81 for restart (Lemacon et al., 2017). The MUS81-cleaved intermediate, a one-ended DSB, may then be repaired by recombination dependent mechanisms such as template switch post-replication repair, which require RAD51, and/or break-induced replication, which requires pol δ, or by translesion synthesis. FANCD2 could participate in such RAD51-mediated fail-safe mechanisms of completing replication (Figure 7). 6) Yet another repair mechanism at stressed replication forks is dependent on post-replication repair of gaps introduced by repriming downstream of lesions on the leading strand, instead of DSBs and could also compensate for or substitute for over-resected reversed forks. In the presence of FANCD2 we see increasing phospho-RPA during stalling which implies activation of ATR, which is presumably necessary for repair. ATR has been shown by fiber tracking to activate PRIMPOL and promote downstream repriming leading to gaps in nascent DNA (Quinet and Vindigni, 2019). In FANCD2-depleted cells, this pathway cannot be activated effectively because RPA-p does not accumulate (Figure 1A, 8 h time point) blunting the checkpoint and recovery and leading to genome instability. FANCD2 has also been reported to counteract NHEJ at IR-induced DSBs, and FANCD2 deficient cells show increased toxic NHEJ, decreased resection, and decreased recombinational repair (Cai et al., 2020). FANCD2 has also been implicated in counteracting Ku70 inhibition of repair (Pace et al., 2010). 7) The strand exchange stimulation function of FANCD2 may be its important contribution to the late stages of ICL repair by the FA pathway, which involves repair of DSBs (Duxin and Walter, 2015). Thus, RAD51 strand exchange stimulation might also explain the minor defect in DSB repair pathways in the absence of FANCD2 reported previously (Nakanishi et al., 2011). The role of FANCD2 in fork protection may be compensated for by either BRCA2, for HR-like fork protection, or in a by-pass mechanism by pol theta/CtIP recruitment for alt-EJ (Kais et al., 2016).

**Figure 7.**
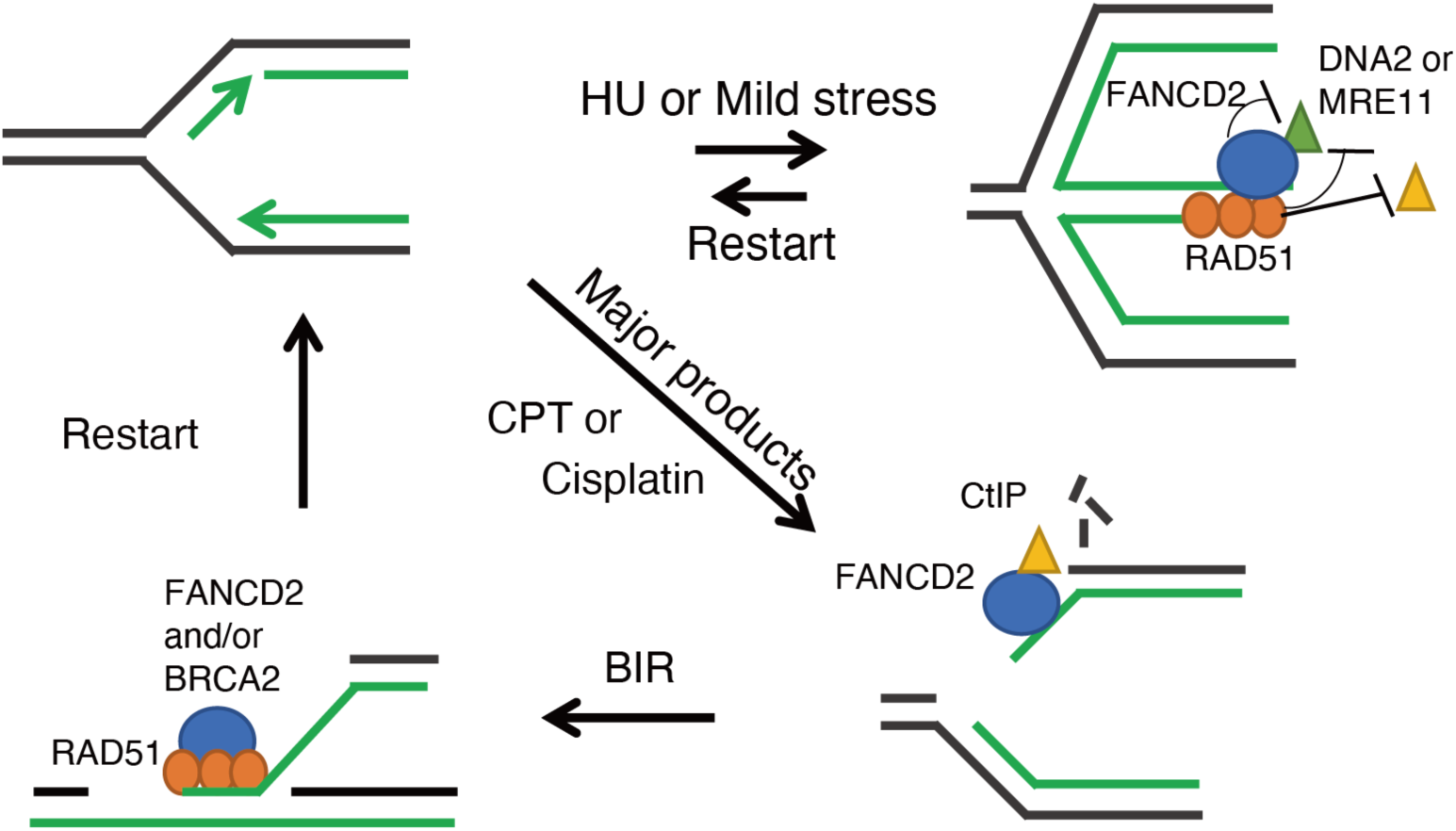
Model for multiple roles of FANCD2 in fork protection studied in this work. At a stalled replication fork upon moderate stress, FANCD2 can protect the regressed arm by directly inhibiting DNA2/MRE11 or stabilizing RAD51 on ssDNA to prevent digestion of nascent DNA by various nucleases. In prolonged stress or at CPT or cisplatin induced damage, FANCD2 can recruit CtIP to the broken fork to facilitate resection and HDR. FANCD2 may also promote RAD51 mediated strand exchange reactions to restart forks either together with BRCA2 or by itself.

### FANCD2 May Act Differently at Different Types of DNA Damage

We observed that while FANCD2 inhibits resection at HU stalled forks, FANCD2 becomes required for repair at more severe types of damage that block forks, such as CPT and cisplatin (Figure 1 and 6). This observation is reminiscent of events at forks stalled by 24 h ICL (interstrand cross-link) inhibition, where FANCD2 recruits CtIP, which augments resection by DNA2/BLM, channeling repair of obligate DSB intermediates in the ICL repair pathway into HR instead of toxic NHEJ, and/or recruits pol theta (Murina et al., 2014; Unno et al., 2014; Yeo et al., 2014).

To recapitulate, our model (Figure 7) taking cumulative data into account, suggests that DNA2 is required for resection of transiently stalled reversed forks to promote restoration of active forks without collapse to DSBs or gaps. FANCD2 is required to keep DNA2/MRE11 mediated resection in a range consistent with preserving genome stability and restoring forks, as demonstrated previously by increased chromosomal aberrations in its absence (Schlacher et al., 2012). However, even if FANCD2 is present, extensive or prolonged stalling, or strong fork blocking lesions, such as CPT or cisplatin, lead to the emergence of genome destabilizing structures that require an additional repair mechanism(s) involving resection. FANCD2 then becomes essential for resection.

### Synthetic Viability and Fanconi Anemia

This study began as a discovery of synthetic viability between DNA2 and FANCD2 defects. We and others have shown that additional FANC alleles show over-resection that is reduced by depletion of DNA2, suggesting that additional proteins are required to fully reconstitute FANCD2-regulated resection at stalled forks (Rickman et al., 2020). Interestingly, different mechanisms of fork protection are seen with different types of damage (Rickman et al., 2020) and with different fork protection factors (Liu et al., 2020). Many additional genes show synthetic viability with FA complementation groups FA-A, FA-C, FA-I, FA-D2, FA, such as BLM helicase (Velimezi et al., 2018). One of the major functions of BLM is to complex with DNA2 in double-strand end resection (Cejka et al., 2010; Nimonkar et al., 2011). BLM is also deleterious for fork protection and replication fork restart in FANCD2 deficient cells, and depletion of BLM rescues restart in that case (Schlacher et al., 2012). It is possible that BLM is also required at reversed forks to promote degradation by DNA2 (and/or EXO1, which is stimulated BLM). These results have significant impact on our goal of using inhibition of DNA2 to increase the therapeutic index for treating Fanconi anemia patients with various cancers by protecting normal, non-cancerous cells from chemotherapeutics that stall replication forks and by limiting inflammation.

## Acknowledgments

This work was supported by a CIHR foundation grant (J.-Y.M.) and J.-Y.M. is a FRQS Chair in genome stability; Korean government grants NRF-2017R1A2B2002289 and RF-2018R1A6A1A0302514 for W.C. sabbatical funding; R50CA211397 to L.Z.; R011CA085344 to B.S.; and USPS grant GM123554 to JLC.

## Author Contributions

Conceptualization, J.C.; Methodology, W.L., I.R., P.P., Y.M., M.C., Y.X., C.L., Q.W., L.Z.; Writing – Original Draft, W.L., P.P., J.C.; Writing – Review and Editing, W.L.,P.P, L.Z., J.M., B.S, J.C.; Funding Acquisition, W.C., J.M., B.S., J.C; Supervision J.M., B.S., J.C.

## STAR Methods

### Key Resources Table

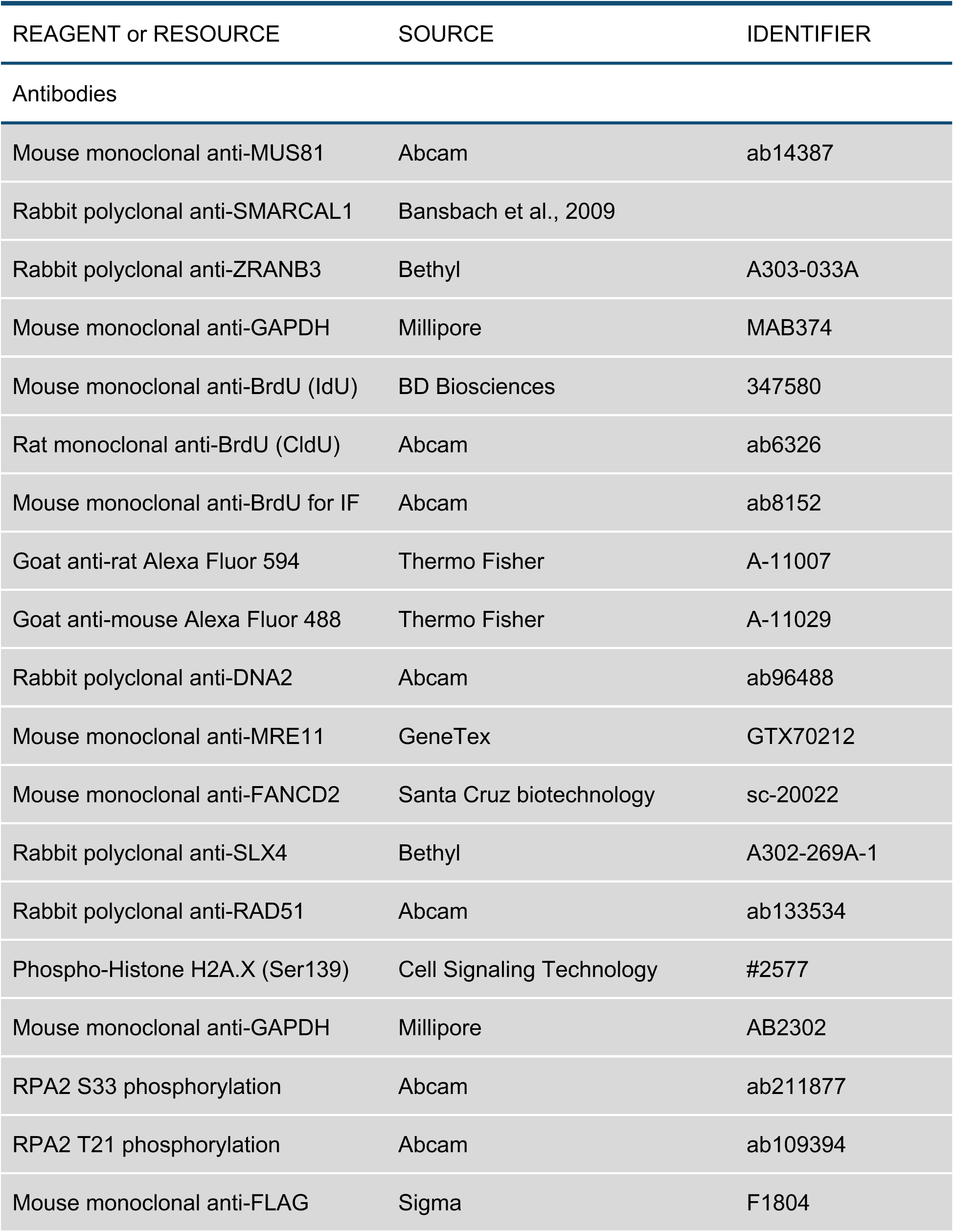
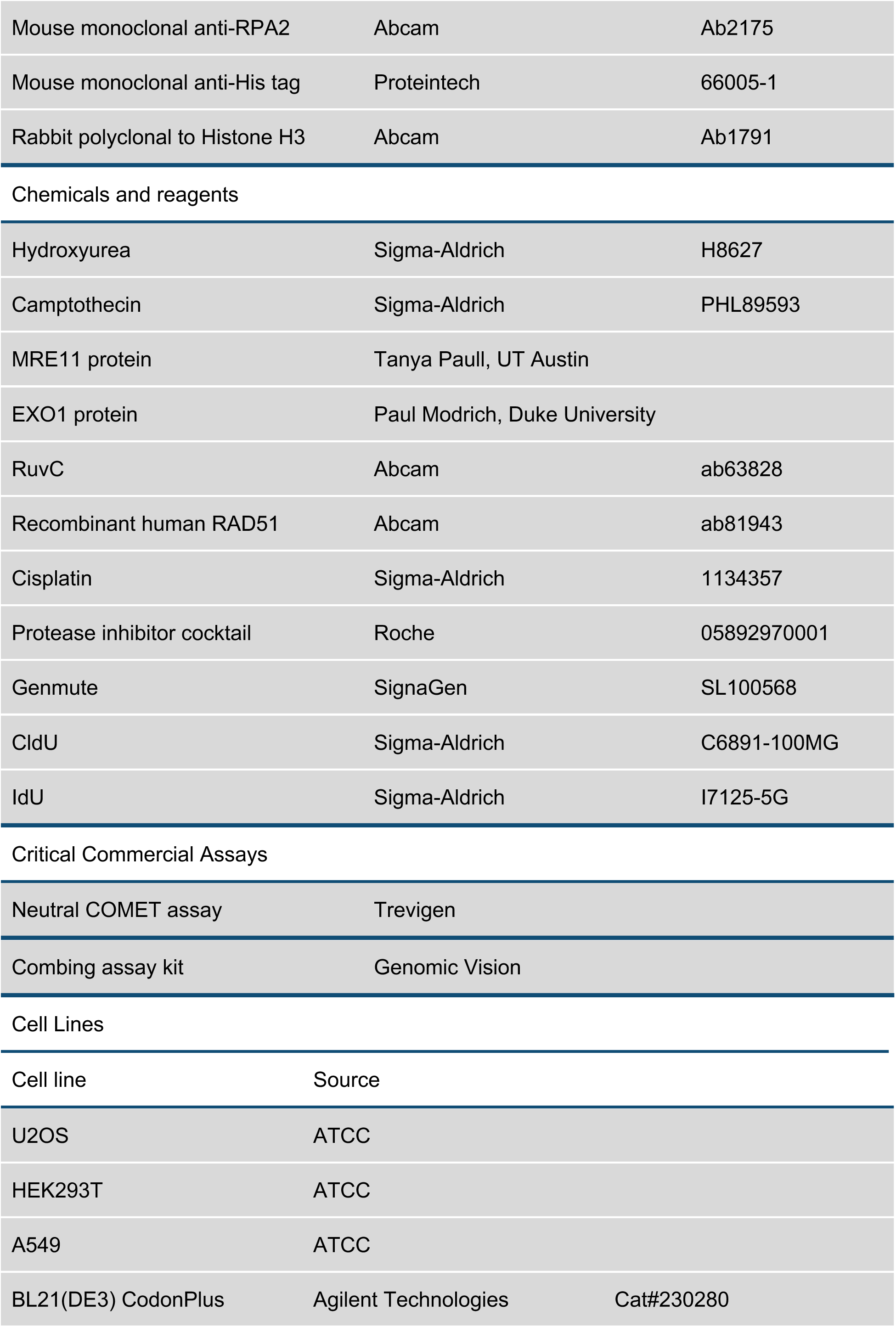
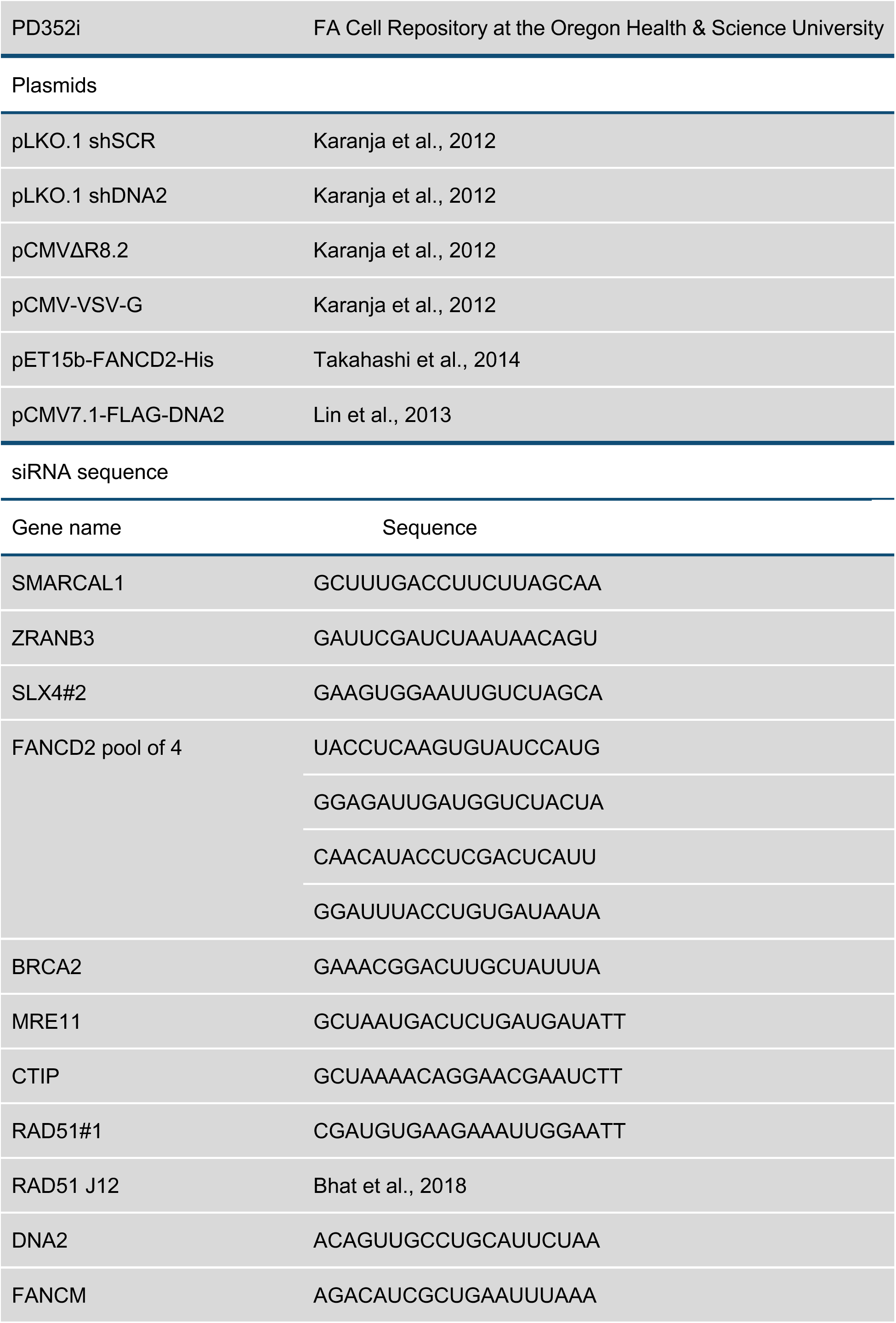
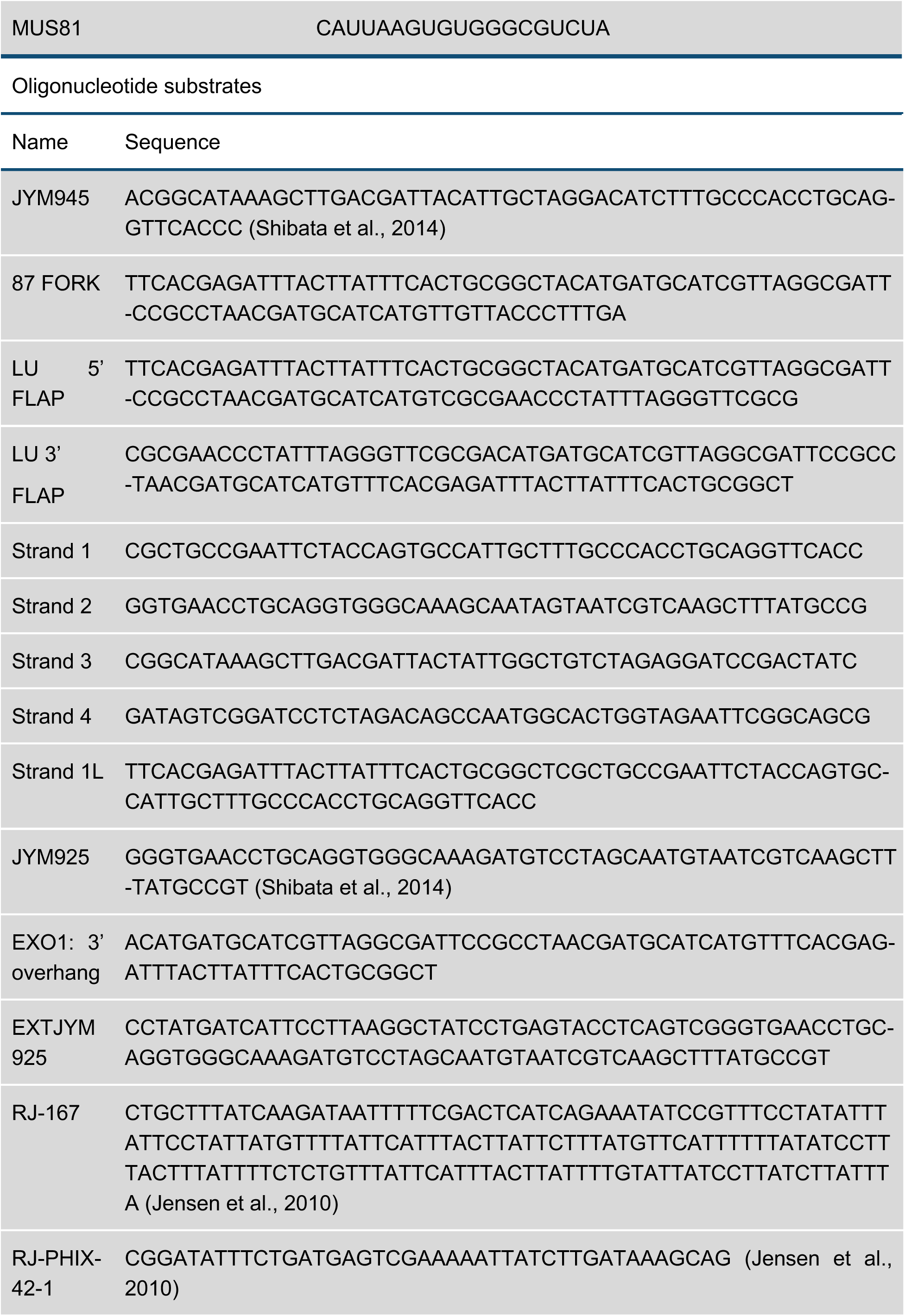
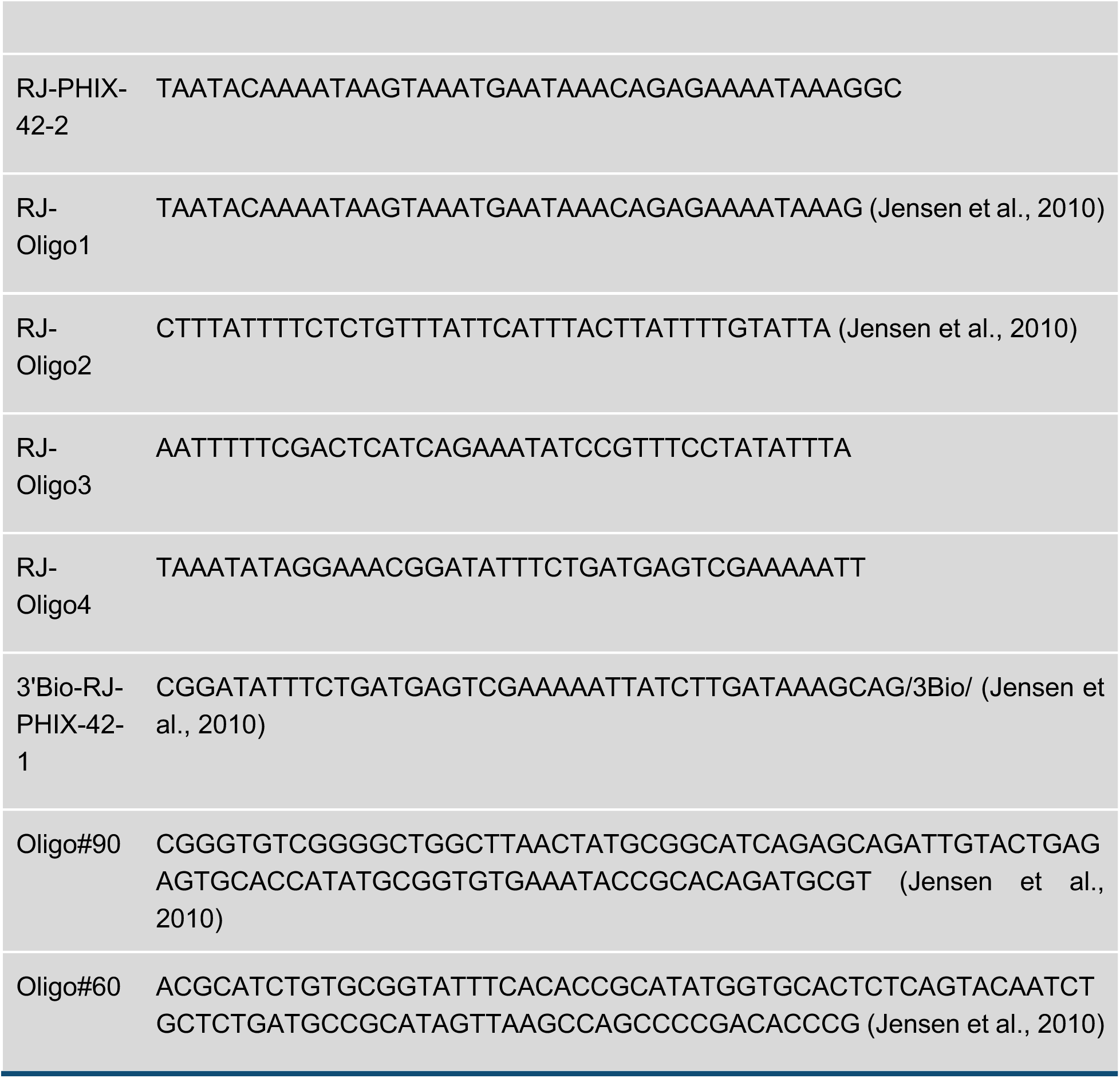

### Lead Contact and Materials Availability

Further information and requests for resources and reagents should be directed to and will be fulfilled by the Lead Contact, Judith L. Campbell. All unique/stable reagents generated in this study are available from the Lead Contact without restriction.

## Method Details

### Cell Culture and Cytotoxicity Assays

U2OS, A549, and PD20 and PD20 with FANCD2 complemented cells were cultured in DMEM medium with 10% FBS.

### Nuclear Fractionation

Cells (1×10^6^) were harvested and washed with PBS, then lysed on ice for 20 min with 100 μl H150 buffer, which contains 50 mM HEPES (pH7.4), 150 mM NaCl, 10% glycerol, 0.5% NP-40 and protease inhibitor cocktail (Roche). The lysate was spun for 10 min at 5000g, and the supernatant is the cytoplasmic fraction. The pellet was washed two times with H150 lysis buffer, and the supernatant discarded. The pellet is the nuclear fraction. The pellet was resuspended in PBS (20 μl) and 20 μl 2xSDS loading buffer and boiled for western blot.

### Immunofluorescence for native BrdU staining and EdU staining

BrdU staining was carried out as described (Couch et al., 2013). Briefly, cells (1×10^5^ labeled with BrdU and EdU as described in the legend to Figure 1 were plated on coverslips, washed with PBS, pre-extracted with ice cold 0.5% Triton-X100 for 4 min, then fixed with 4% paraformaldehyde for 10 min, permeabilized with 0.1% Triton-X100 for 2 min and then washed with PBS 3 times. Blocking was carried out with 1% BSA in PBS for 1 h. For EdU detection, a 1 ml click reaction containing 5 μl 1 mM Azide-488 (Invitrogen), 100 μl 20 mg/ml sodium ascorbate, 20ul 100mM CuSO_4_) was performed to detect incorporated EdU. Then FANCD2 antibody 1:200 in blocking buffer incubated overnight at 4 degree. For BrdU staining, slides were incubated with BrdU and FANCD2 primary antibody overnight at 4°C. The slides were washed in PBS three times and then incubated with secondary antibody (1:200, Alexa Fluor 594 and 488 from Invitrogen) for 1 h at room temperature. The slides were washed with PBS 3 times and mounted with Prolong Gold AntiFade Reagent with DAPI (Invitrogen P36941).

### Plasmid and siRNA Transfection

A549 and U2OS cells were plated the day before transfection. 20 nM siRNA was used for single and 16 nM for each siRNA in co-transfection. Cells was transfected with Genmute and labeled as indicated 72 hours post-transfection. DNA2 plasmid transfection was described previously (Liu et al., 2016).

### DNA Fiber Assay

DNA fiber spreading and staining were performed as previously described (Liu et al., 2016). Briefly, 1000 labeled cells (2 μl, 500 cells per μl) on slides were half dried, 10 μl lysis buffer (0.5% SDS, 200 mM Tris-HCl pH 7.4, 50 mM EDTA) was added, followed by incubation for 6 min at room temperature. The slide was tilted to 15 degrees to allow the DNA to run slowly down the slide. Slides were air dried for at least 40 minutes, and fixed for 2 min in 3:1 methanol: acetic acid in a coplin jar. Slides were dried in a hood for 20 min. Slides were treated with 2.5 M HCl for 70 min for denaturation and then washed with PBS 3 times and blocked with 10% goat serum in PBST (0.1% Triton-X100 in PBS) for 1 h. Slides were incubated with the rat anti-BrdU and mouse anti-BrdU antibody, 1:100, for 2 h, washed 3 times with PBS, and then incubated with secondary antibody (Goat anti-Mouse 488 and Goat anti-Rat 594, Invitrogen) at 1:200. Slides were imaged with immunofluorescence microscopy and fiber length measured by Nikon software. Statistical analyses were completed using Prism. An ANOVA test was used when comparing more than two groups followed by a Dunnett multiple comparison post-test.

### Neutral COMET assay

The neutral COMET assays were performed in accordance with the manufacturer’s (Trevigen) instructions. Cells were trypsinized and washed, then palleted, resuspended with low melt agarose, then dropped on the slides. After cool down, the slides were incubated in cold lysis buffer (Trevigen) for 1 hour, then incubated in running buffer for 30 minutes, and then subjected to electrophoresis at 21 volts for 45 minutes. Slides were then immersed in precipitation buffer (Trevigen) and 70% ethanol for 30 minutes, respectively. Slides were dried overnight and stained with SYBR green I (Thermofisher). Slides were imaged with fluorescence microscope with FITC channel.

### Immunoprecipitation

For FLAG pulldown assays and immunoprecipitation assays, 293T cells were transfected with or without RAD51 vector (or FLAG-DNA2 vector) using the Polyjet (SignaGen SL100688) transfection reagent. 24 h after transfection, the cells were incubated with or without 2 mM HU for 3 h. Cells (1×10^7^) were collected and lysed by brief sonication and incubation in the immunoprecipitation (IP) buffer H150 (50 mM HEPES-KOH (pH7.4), 150 mM NaCl, 0.1% NP40 and 10% glycerol) with protein inhibitor cocktail (Thermo Fisher) for 30 min. After centrifugation (20,000g, 15 min, 4°C), the supernatants were collected, and the protein concentration determined. Cell lysate (1mg) was pre-cleaned with 10 μl Protein A/G beads (Thermo #88802) for 1 h. After removing beads, the lysate was incubated with 2 μg (1μg/μl) anti-RAD51 (ab133534 Abcam) or anti-FLAG M2 magnetic beads for FLAG pulldown (Sigma). Then 10 μl Protein A/G magnetic beads were added and incubated overnight at 4°C. The beads were washed three times with the IP buffer H150 and boiled in 1x SDS-PAGE loading buffer directly. The DNA2 and FANCD2 were analyzed by western blot analysis.

### Oligonucleotides

Oligonucleotide substrates for enzymatic assays were labeled at the 5’ end with ^32^P using polynucleotide kinase. The sequences are listed in the Key Resources Table. For DNA2 assays, single-stranded DNA was JYM945 (Shibata et al., 2014). The forked substrate was designated 87 FORK. The 5’ flap substrate was LU 5’ FLAP. The 3’ flap substrate was LU 3’. The reversed fork with blunt ends consisted of 4 oligonucleotides: strand 1, strand 2, strand 3 and strand 4 in the Key Resources Table (van Gool et al., 1998). The reversed fork with 5’ overhang consisted of strand1L, strandFANCD2, strand3, and strand4.

The MRE11 nuclease duplex substrate was formed by annealing 5’ labeled JYM945 to JYM925 (Shibata et al., 2014). This was also used for binding of RAD51 to dsDNA. For the EXO1 assay, a hairpin with a 3’ overhang was used.

For RAD51 binding, JYM945 was used. For RAD51 strand exchange assays the single-stranded DNA was EXTJYM925: The 60mer duplex was formed by annealing labeled JM945 to JM925 (see MRE11 substrate).

### Proteins

Recombinant human RAD51 was from Abcam (ab81943) and tested for ATPase, strand exchange, and DNA binding. RuvC was Abcam (ab63828). MRE11 was the gift of Tanya Paull (UT Austin) and EXO1 (0.77 mg/ml) was a gift from Paul Modrich, Duke University. Sources of FANCD2 and DNA2 are described in the text or figure legend describing the experiments in which they were used.

### FANCD2-His Purification from *E. coli*

Human FANCD2 protein was purified from *E. coli* as previously described (Takahashi et al., 2014). The FANCD2 vector was transformed into BL21(DE3) CodonPlus (Agilent Technologies 230280) cells. 20 L of transformed cells were amplified at 30°C, 250 rpm. FANCD2 protein was produced by adding 0.5 mM IPTG at 16°C for 18 hours, when the cell density reached an OD_600_=0.6. The *E. coli* cells were harvested and pelleted and lysed in Buffer A (50 mM Tris-HCl PH8.0, 500 mM NaCl, 5 mM 2-mercaptoethanol, 1 mM phenylmethylsulfonyl fluoride (PMSF), 12 mM imidazole, and 10% glycerol), and disrupted by sonication. The lysate was centrifuged at 20,000 g at 4°C; the supernatant was mixed gently by the batch method with 3ml of Ni-NTA agarose beads, at 4°C for 1h. The beads were packed into an Econo-column, and were washed with 67 column volumes of buffer A. The His-tagged FANCD2 were eluted with a 20 column volumes linear gradient of 12-400 mM imidazole in buffer A. The peak fractions were collected. To remove His tag from the FANCD2 protein, thrombin protease (2U/mg GE healthcare) was added, and the sample was then dialyzed against 4L of buffer B (20 mM Tris-HCl, pH8.0, 200 mM NaCl, 5 mM 2-mercaptoethanol, 10% glycerol). Afterward, the sample was passed through a Q Sepharose Fast Flow (2.5 ml, GE Healthcare) column. The resin was washed with 60 column volumes of buffer B containing 250 mM NaCl. Human FANCD2 was then eluted with a 20-column volume linear-gradient of 250 mM-450 mM NaCl in buffer B. The peak fractions were collected, and human FANCD2 was further purified by gel filtration chromatography on a Superdex 200 column (GE Healthcare) equilibrated with Buffer B containing 200 mM NaCl. The purified FANCD2 was concentrated, frozen in aliquots, and stored at −80°C. The concentration of purified FANCD2 was determined by the Bradford method, using BSA as standard.

### FLAG-DNA2 Purification from Mammalian cells

The FLAG-DNA2 expression and purification procedure was as described previously (Lin et al., 2013). In brief, whole cell lysates were incubated with the M2 FLAG magnetic beads (Sigma) for at least 6 h in cold room. After extensively washing with a buffer containing 50 mM Tris-Cl (pH 7.5) and 500 mM NaCl, the bound proteins were eluted with 3X FLAG peptide (Sigma). The purity of DNA2 proteins was analyzed by 4–15% gradient SDS–polyacrylamide electrophoresis (SDS–PAGE) and Coomassie brilliant blue staining, and the concentration was determined by comparison to BSA after Coomassie blue staining of SDS gels.

### Mapping the FANCD2 binding domain in DNA2

Mutant FLAG-DNA2 proteins were prepared using site-directed mutagenesis. The N-terminal deletions were made using the HiFi DNA cloning kit from NEB to excise portions of the N-terminus of the gene, while C-terminal deletions were made by the insertion of a stop codon earlier in the gene construct. Coimmunoprecipitations were performed by overexpressing the DNA2 proteins in HEK-293T cells prior to making cell lysates. FANCD2 was added to the lysates to a final concentration of 2 nM protein to ensure measurable interaction with DNA2. The FANCD2:DNA2 complex was pulled down using a FANCD2 antibody attached to magnetic beads. The beads were washed prior to eluting the samples using SDS loading buffer, and the samples were analyzed by western blot using a 3XFLAG antibody.

### Strand Exchange Assays

Single-stranded DNA (EXTJYM925) was preincubated in the presence of RAD51 and FANCD2 in a reaction mixture containing 25 mM TrisOAc (pH 7.5), 2 mM MgCl2, 2 mM CaCl2, 2 mM ATP, 1 mM DTT and 0.1 mg/ml BSA for 5 min at 37°C for filament formation. Following pre-incubation, dsDNA (5’ labeled JYM945 annealed to JYM925) with the labeled strand complementary to the filament, was added to the reaction mixture and incubation was continued for an additional 30 min at 37°C for strand exchange. Reactions were terminated by the addition of proteinase K and SDS to 0.5 mg/ml and 0.25% respectively and incubated for 10 min at 37°C. 1 µL of Loading Buffer (2.5% Ficoll-400, 10 mM Tris-HCl, pH 7.5, and 0.0025% xylene cyanol) was added and samples were loaded on an 8% native gel using 29:1 30% acrylamide solution. Gels were run at 100v (constant voltage) for 4 h.

For strand exchange assays that used the 3’ overhang DNA (RJ-167 annealed to RJ-PHIX-42-1) for filament formation during preincubation, 5’ labeled dsDNA (5’ labeled RJ-Oligo1 annealed to RJ-Oligo2) was used as its respective strand exchange target during the 30 min incubation. Similarly, in instances using 5’ overhang DNA (RJ-167 annealed to RJ-PHIX-42-2) to generate filaments, 5’ labeled dsDNA (5’ labeled RJ-Oligo4 annealed to RJ-Oligo3) was used as its double-stranded target.

### Biotin Pull-down assays for RAD51 and FANCD2 association with overhang DNA

The protocol was adopted from Jensen et al., 2010. Briefly, the oligonucleotide substrate Bio-RJ-PHIX-42-1 composed of the same sequence as RJ-PHIX-42-1 but containing a 3′ biotin modification was obtained from IDT (Integrated DNA Technologies) and PAGE purified. The biotinylated 3’ overhang substrate was generated by annealing Bio-RJ-PHIX-42-1 to oligonucleotide RJ-167 at a 1:1 molar ratio in STE buffer. Competitor heterologous dsDNA was similarly generated by annealing PAGE purified oligonucleotides Oligo#90 and Oligo#60. For pull-down, RAD51 and FANCD2 proteins were incubated in Buffer S (25 mM TrisOAC pH 7.5, 1 mM MgCl2, 2 mM ATP, 1 mM DTT, and 0.1 µg/µL BSA) for 15 min at 37°C followed by the addition of 3’ overhang DNA (162 nt RJ-167 annealed to 42 nt 3’ Bio-RJ-PHIX-42-1) and competitor heterologous dsDNA (90mer, Oligo #90/Oligo #60 oligonucleotides) and the reaction was incubated for an additional 5 min at 37°C. Where DNA was omitted, TE buffer was used and similarly, respective proteins storage buffers were used where proteins were omitted. DNA-protein complexes were captured by adding the reaction mixtures to 2.5 µL of MagnaLink Streptavidin magnetic beads (Solulink) pre-washed by excess Buffer S supplemented with 0.1% Ipegal CA-630 and rotating for 10 min at 25°C. Bead complexes were then washed with excess Buffer S supplemented with 0.1% Ipegal CA-630. Protein was then eluted by re-suspending in 15 µL of 2X protein sample buffer and heating at 54°C for 4 min. The elution fraction was then loaded into a Bis-Tris protein gel for western analysis. Following transfer, the membrane was cut horizontally at the 70 kD marker to separately probe for RAD51 and FANCD2. The lower half was probed using 1:1000 diluted α-RAD51 (Abcam) and the upper half using 1:1000 diluted α-FLAG (ThermoFisher) to detect FANCD2. Anti-mouse (LI-COR) secondary antibody diluted 1:10,000 was used and membranes were imaged via Odyssey imaging system. Bands were quantified using ImageQuant (Cytiva) software.

### Nuclease and DNA-dependent ATPase Assays

#### DNA2 nuclease assay

FANCD2-His or FANCD2-His diluent was incubated in DNA2 nuclease reaction mix (50 mM HEPES-KOH, pH 7.5, 5 mM MgCl_2,_ 2mM DTT, 0.25 mg/ml BSA) for 30 min at 4°C. DNA2, preincubated with substrate (87 fork, 1.5 nM molecules) for 5 min on ice, was added and the reaction was incubated for 30 min at 37°C. See Key Resources Table for substrate sequences. Following incubation, proteinase K and SDS were added to 1 mg/ml and 0.5%, respectively, and incubation continued for 10 min at 37°C. Denaturing termination dye (2X: 95% deionized formamide, 10 mM EDTA, 0.1% bromophenol blue and 0.1% xylene cyanol) was added and the mixture boiled for 5 min. Samples were run on a sequencing gel and the gel analyzed by phosphor imaging.

#### MRE11 nuclease assay

MRE11 reaction mixtures contained 25 mM MOPS (pH 7.0), 60 mM KCl, 0.2% Tween 20, 2 mM DTT, and 1 mM MnCl_2_ as described (Paull and Gellert, 1998). MRE11 and blunt dsDNA 60mer substrate (JYM925/JYM945 oligonucleotides) were incubated together on ice for 5 min before being introduced to the reaction mixture at 200 nM and 1 nM, respectively, and incubated for 30 min at 37°C. Following incubation, reactions were terminated by adding proteinase K and SDS was added to 1 mg/ml and 0.5% respectively and incubated for 10 min at 37°C. 10 µl of 2X termination dye was added and samples boiled for 5 min. After denaturing, samples were run on a 12% sequencing gel at constant 60W and the gel analyzed by phosphor imaging.

#### EXO1 nuclease assay

Conditions are as previously described (Shao et al., 2014). Reaction mixtures (10 µl) contained 20 mM Tris-HCl, pH 7.6, 0.75 mM HEPES-KOH, 120 mM KCl, 250 μg/ml BSA, 2 mM ATP, 1 mM glutathione, 2 mM MgCl_2_, 1% glycerol, 0.06 mM DTT, 1.5 nM substrate and EXO1 (0.77 nM).

#### ATPase assays

The DNA-dependent ATPase assays were carried out as previously described (Masuda-Sasa et al., 2006). Reaction mixtures (10 µL) contained 20 mM TrisOAc (pH 7.5), 4 mM MgCl2, 4 mM CaCl2 (where shown), 1 mM DTT, 0.5 mM ATP, 20 µCi/ml [γ-32P]-ATP, and 900 nM (in nucleotides) of cold ssDNA (60 nt, oligonucleotide JYM945). Indicated concentrations of FANCD2 and 300 nM RAD51 were incubated in reaction mixture for 90 min at 37°C and then reactions were stopped by the addition of EDTA to 4 mM. All reactions contained equal amounts of FANCD2 diluent.

### Quantification and Statistical Analysis

Statistical analyses were completed using Prism. An ANOVA test was used when comparing more than two groups followed by a Dunnett multiple comparison post-test. A two-tailed t-test was used to compare two samples with normally distributed data. No statistical methods or criteria were used to estimate sample size or to include/exclude samples.

### Data and Code Availability

