## Supplemental figures 1-3 for "FANCD2 and RAD51 recombinase directly inhibit DNA2 nuclease at stalled replication forks and FANCD2 acts as a novel RAD51 mediator in strand exchange to promote genome stability"

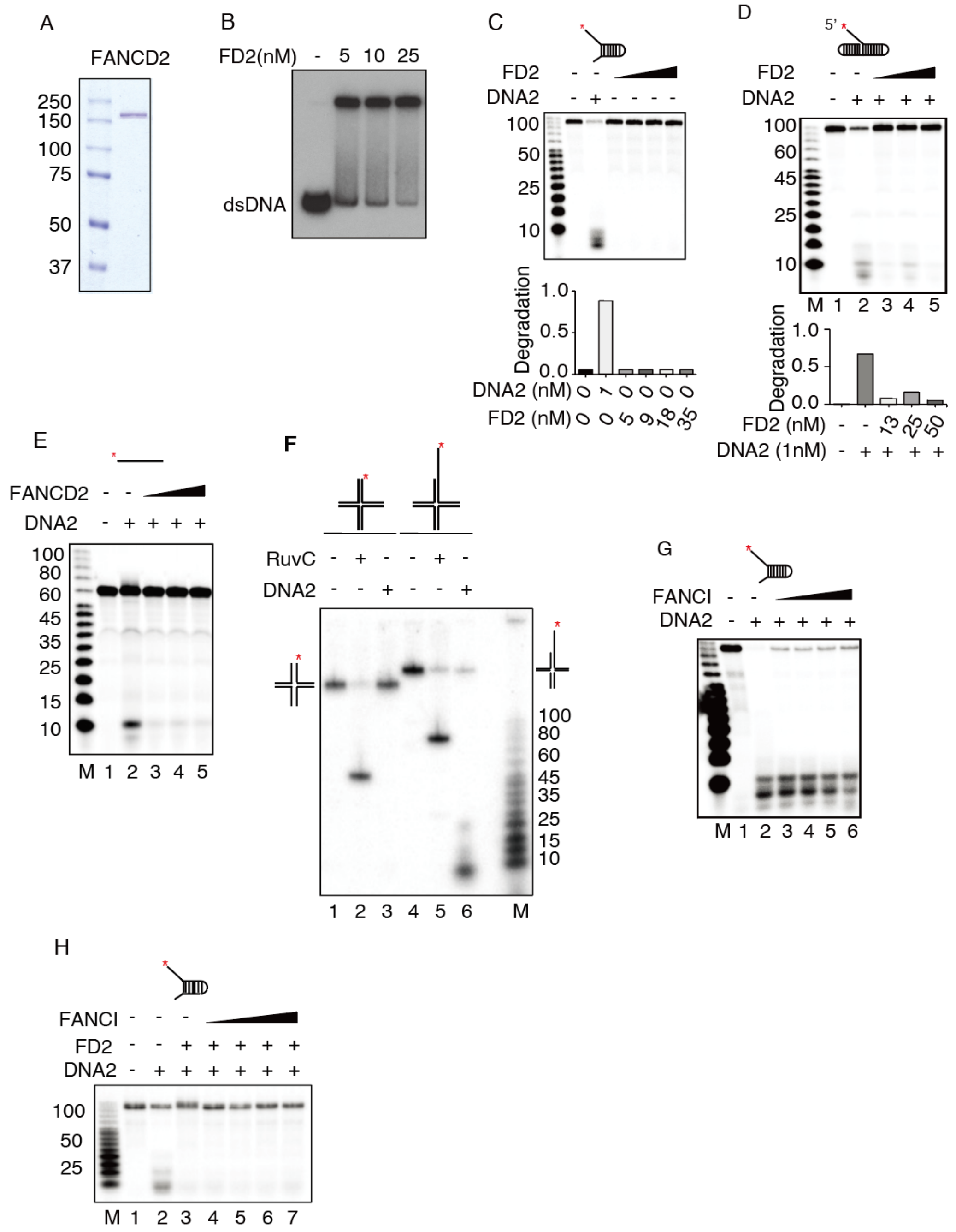
A
FANCD2
250
150
100
75
50
37
B
FD2(nM) - 5 10 25
dsDNA
C
FD2
DNA2
100
50
25
10
Degradation
1.0
0.5
0.0
DNA2 (nM) 0 0 1 0 0 0 0
FD2 (nM) 0 0 0 5 9 18 35
D
5'
FD2
DNA2
100
60
45
25
10
M 1 2 3 4 5
Degradation
1.0
0.5
0.0
FD2 (nM) - - 13 25 50
DNA2 (1nM) - + + + +
E
FANCD2
DNA2
100
80
60
45
35
25
15
10
M 1 2 3 4 5
F
RuvC
DNA2
100
80
60
45
35
25
15
10
1 2 3 4 5 6 M
G
FANCI
DNA2
M 1 2 3 4 5 6
H
FANCI
FD2
DNA2
100
50
25
M 1 2 3 4 5 6 7

**Supplementary Figure 1 (related to Figure 1). FANCD2-mediated Inhibition of in vivo and in vitro resection by DNA2**

(A) Human FANCD2-His was expressed and purified from insect cells as described (Roques et al., 2009). Gel analysis of the protein is shown.

(B) dsDNA binding by the FANCD2-His was carried out as described (Niraj et al., 2017).

(C) FANCD2 does not show nuclease activity under the conditions of DNA2 nuclease assay, as described in Method Details. Markers show specific DNA2 nuclease product at 10 nt and smaller. Lane 1, DNA forked substrate without protein; lane 2, DNA2 alone; lanes 3-6, FANCD2 alone, the concentration is 5, 9, 18, 35 nM.

(D) FANCD2 inihibits DNA2 on a flap substrate. The assay was conducted as in Figure 1C, FANCD2 concentrations are 13, 25, 50 nM, respectively.

(E) FANCD2 inhibits DNA2 on ssDNA. with JYM945 substrate DNA (1.5 nM), FANCD2 concentrations are 13, 25,50nM, respectively.

(F) Oligonucleotides described in Supplementary Table S2 were labeled, annealed and gel purified. The reversed fork with blunt ends consisted of 4 oligonucleotides: 5' labeled strand 1, strand 2, strand 3, and strand 4 (van Gool et al., 1998). The reversed fork with 5' overhang consisted of 5' labeled strand 1L: strand 2, strand 3, and strand 4. 1.5 nM of the indicated substrate, 5' end-labeled as indicated, was incubated in a 10 μl reaction mix containing 25 nM RuvC (Abcam) or diluent, 50 mM Tris-HCl (pH 8.0), 10 mM MgCl2, 100 μg/ mL BSA,1 mM DTT at 37° C for 30 min (Amunugama et al., 2018). Following incubations, proteinase K and SDS were added to 1 mg/ml and 0.5% respectively and incubated for 10 min at 37° C. 1 μl of

10X native loading dye was added to final 2.5% Ficoll- 400, 10 mM Tris-HCl (pH 7.5) and 0.0025% xylene cyanol concentrations. Samples were separated on an 8% native gel using 29:1 30% acrylamide solution, constant voltage 200V in cold room, 1X TAE, 120 min, 5 h exposure. The labeled product on the blunt ended reversed fork was expected to be 50 bp and 80 bp on the recessed reversed fork.

(G) FANCI does not inhibit DNA2. Reaction conditions were as in Figure 1C and FANCI was added at the concentrations indicated at the same time as DNA2.

(H) FANCI does not stimulate inhibition of DNA2 by FANCD2. Reaction conditions were as in Figure 1C and FANCI was added at the concentrations indicated at the same time as DNA2. FANCI concentrations are 8, 16, 32, 75 nM, respectively.

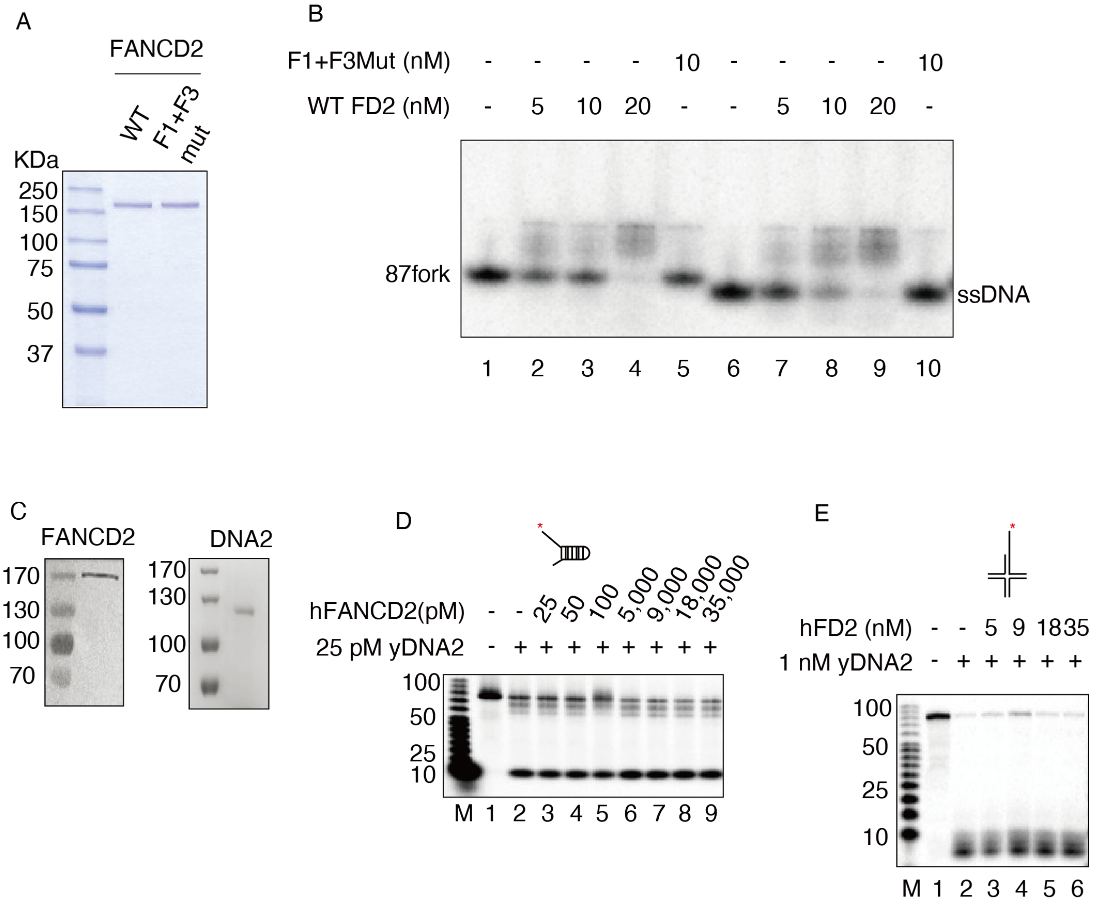


**Supplementary Figure 2. Related to Figure 2. On the roles of FANCD2/DNA and FANCD2/DNA2 protein/protein reaction in the inhibition of DNA2 by FANCD2.**

(A) Human FANCD2 WT and FANCD2-F1+F3 Mut-His purification gel analysis.

(B) Comparison of DNA binding by FANCD2 and FANCD2-F1+F3Mut. Reaction conditions were as in (Niraj et al., 2017). Reactions in lanes 1-5 contained 1 nM 5' labeled 87FORK and reactions in lanes 6-10 contained 1 nM ssDNA: 5' labeled JYM945* (60nt). Imagequant quantification revealed a 10- fold reduction in FANCD2-F1+F3Mut binding to ssDNA (lane8 vs lane 1).

(C) Purified FANCD2-His and FLAG-DNA2. Proteins were purified as described in Method Details and analyzed by SDS gel.

(D) Yeast DNA2 is not inhibited by FANCD2 – 87 FORK substrate. Conditions were the same as in Figure 1C except that yeast FLAG- DNA2 (Masuda-Sasa et al., 2006) was used instead of human. Note that yeast DNA2 is more active than human DNA2 and therefore the amounts of yeast DNA2 shown are adjusted to give ≤ 50% degradation of substrate.

(E) Yeast DNA2 is not inhibited by FANCD2 – reversed fork substrate. Conditions were the same as in Figure 1 C except that yeast FLAG-DNA2 (Masuda-Sasa et al., 2006) was used instead of human DNA2.

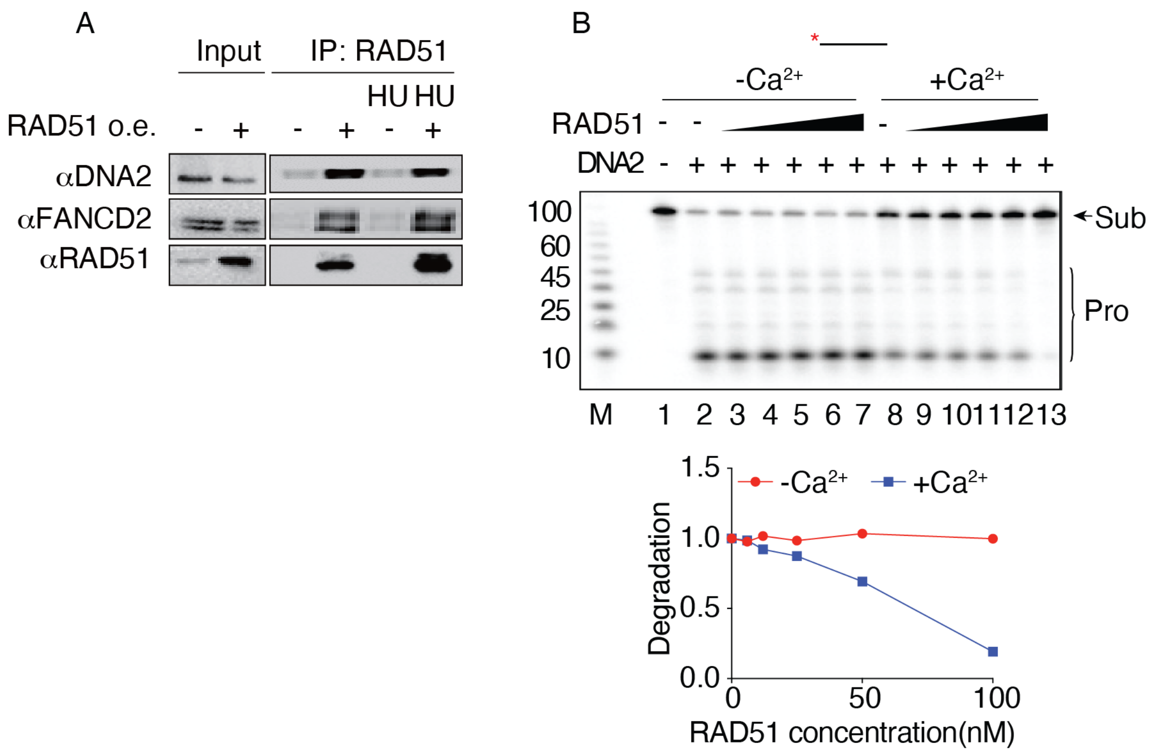


**Supplementary Figure 3 (Related to Figure 3). RAD51 filaments inhibit DNA2.**

(A) Co-immunoprecipitation of FANCD2 and RAD51 using RAD51 antibody for immunoprecipitation. RAD51 vector or empty vector was transfected into 293T cells and cells were split 24 hours post-transfection. At 48 hours post-transfection, cells were incubated with or without 2 mM HU for 3 hours; cells were then harvested and lysed. Lysates were incubated with RAD51 antibody and IgG-agarose beads, then washed with lysis buffer. Immunoprecipitates were analyzed for FANCD2 and DNA2 by western blotting on an 8% acrylamide gel.

(B) RAD51 filaments inhibit DNA2 on a ssDNA substrate. Increasing amounts of RAD51 were preincubated with 4 nM ssDNA (JYM945) in 25 mM TrisOAc (pH 7.5), 2 mM $MgCl_2$, 2 mM ATP, 0.1 mg/ml BSA, and 2 mM DTT for 10 min at 37°C, conditions shown to be permissive for both filament formation and DNA2 nuclease activity in control assays (not shown). DNA2 was then added to 5 nM as indicated and reactions incubated for 30 min at 37°C. Samples were processed, run on a sequencing gel and analyzed by image J. $Ca^{2+}$ (2 mM) was present in lanes 8-13.
